# The human mitochondrial mRNA structurome reveals mechanisms of gene expression

**DOI:** 10.1101/2023.10.31.564750

**Authors:** J. Conor Moran, Amir Brivanlou, Michele Brischigliaro, Flavia Fontanesi, Silvi Rouskin, Antoni Barrientos

**Affiliations:** Department of Biochemistry and Molecular Biology. University of Miami Miller School of Medicine. 1600 NW 10th Ave. Miami, FL-33136 (USA); Department of Microbiology. Harvard Medical School. 77 Ave. Louis Pasteur. Boston, MA-02115 (USA); Department of Neurology. University of Miami Miller School of Medicine. 1600 NW 10th Ave. Miami, FL-33136 (USA); The Miami Veterans Affairs (VA) Medical System. 1201 NW 16th St, Miami, FL-33125 (USA)

**Keywords:** Mitochondrial mRNA structurome, mitoDMS-MaPseq, programmed ribosome frameshifting, DREEM, mitochondrial gene expression, RNA secondary structure

## Abstract

The mammalian mitochondrial genome encodes thirteen oxidative phosphorylation system proteins, crucial in aerobic energy transduction. These proteins are translated from 9 monocistronic and 2 bicistronic transcripts, whose native structures remain unexplored, leaving fundamental molecular determinants of mitochondrial gene expression unknown. To address this gap, we developed a mitoDMS-MaPseq approach and used DREEM clustering to resolve the native human mitochondrial mt-mRNA structurome. We gained insights into mt-mRNA biology and translation regulatory mechanisms, including a unique programmed ribosomal frameshifting for the *ATP8/ATP6* transcript. Furthermore, absence of the mt-mRNA maintenance factor LRPPRC led to a mitochondrial transcriptome structured differently, with specific mRNA regions exhibiting increased or decreased structuredness. This highlights the role of LRPPRC in maintaining mRNA folding to promote mt-mRNA stabilization and efficient translation. In conclusion, our mt-mRNA folding maps reveal novel mitochondrial gene expression mechanisms, serving as a detailed reference and tool for studying them in different physiological and pathological contexts.

## Introduction

Expression of the human mitochondrial genome (mtDNA) provides thirteen essential protein subunits of the oxidative phosphorylation (OXPHOS) enzymatic complexes that catalyze aerobic energy transduction and support life. The mtDNA also encodes the 2 ribosomal RNAs (12*S* and 16*S* rRNA) and 22 transfer RNAs (tRNAs) required for synthesizing these proteins in mitochondrial ribosomes (mitoribosomes) ^1^. Over the past decades, the molecular machineries and mechanisms governing mtDNA transcription and mitochondrial messenger RNA (mt-mRNA) stability, processing, modification, and translation have been progressively characterized ^1–9^, and remain the subject of intense investigations. However, our knowledge of the mt-mRNA folding landscape or structurome is very limited, which has hindered our mechanistic understanding of mtDNA gene expression and its regulation.

The human mtDNA is a compact double-stranded 16.569 Kb molecule consisting of heavy (H) and complementary light (L) strands. Transcription spans almost their entire length, producing two long polycistronic transcripts. The H-strand primary transcript contains mRNAs for 12 OXPHOS subunits, 2 rRNAs (12*S* and 16*S*), and 14 tRNAs ^10^. The L-strand primary transcript contains 1 mRNA that encodes subunit 6 of NADH dehydrogenase (ND6), 8 tRNAs, and non-coding RNAs (ncRNAs) like 7S ncRNA that regulates transcription initiation ^11^. In both strands, the rRNAs and most mRNAs are flanked by tRNAs, whose processing by endoribonuclease (RNase) P and RNase Z (ELAC2) yields the individual transcripts ^12–14^. Processing of non-canonical junctions lacking tRNAs (at the 3′ UTR of *ND6*, the 5′ UTR of *COX1,* the *COB-ND5* junction, and the *ATP8/6-COX3* junction) is regulated by members of the FASTKD protein family ^15^. Among the 11 mature mRNAs, 2 transcription units remain unprocessed as bicistronic elements containing overlapping and −1 shifted open reading frames encoding ATP8 and ATP6, or ND4L and ND4. Mitochondrial mRNAs (except *ND6*) are post-transcriptionally polyadenylated at their 3’ end ^16^, which both completes the UAA stop codon of a majority of mRNAs and influences their stability ^17,18^. These mRNAs lack Shine-Dalgarno sequences for ribosomal recognition as in prokaryotes, and have very short or no 5’ and 3’ untranslated regions ^19^. The mechanism for the mitoribosome locating start codons and initiating translation of these leaderless mRNAs is not fully understood. Early *in vitro* studies showed preferential selection of the 5’ terminal start codon on leaderless mRNAs over an internal AUG in mammalian mitoribosomes ^20^. Recent cryo-EM reconstructions suggest that a complex formed by the Leucine-rich pentatricopeptide-repeat-containing (LRPPRC) protein and stem-loop RNA-binding protein (SLIRP), known to contribute to RNA stability and polyadenylation ^21–23^, also operates in the targeting of mRNAs to the mitoribosome ^7,24^. Pioneering *in vitro* mRNA structure determination of the first 35 nucleotides for all bovine mitochondrial mRNAs using Selective 2′ Hydroxyl Acylation analyzed by Primer Extension (SHAPE) chemistry showed that they are highly unstructured, single-stranded, and thus potentially poised for ribosome binding ^25^. However, obtaining mRNA structure data within intact mitochondria is critical to the understanding of its biological context.

Recently, several methods and chemical reagents have emerged for probing RNA structures in living cells. However, these techniques have not yet been applied to unravel the folding patterns of mammalian mitochondrial transcripts. These methods utilize cell-permeant chemical reagents to modify accessible nucleotides when they are in single-stranded RNAs ^26^. One such chemical probe is dimethyl sulfate (DMS), which methylates the base-pairing faces of unpaired and accessible adenines and cytosines ^27^. DMS mutational profiling with sequencing (DMS-MaPseq) is a high-throughput, genome-wide RNA structure probing strategy that takes advantage of a high-fidelity and processive thermostable group II reverse transcriptase (TGIRT) enzyme that reads DMS modifications as mismatches ^28–30^. The approach captures the dynamic nature of RNA folding, which is then revealed by the clustering algorithm DREEM (Detection of RNA folding Ensembles using Expectation-Maximization) ^31^, identifying coexisting alternative conformations that are adopted by the same RNA sequence based on DMS-MaPseq data.

Here, we have optimized and employed a mitochondrion-specific approach, mitoDMS-MaPseq, to investigate the structure of mt-RNAs in functional, isolated human mitochondria. By using DREEM for data analysis, we achieved an accurate and high-confidence understanding of the mt-RNA folding landscape. This information, in combination with *in organello* metabolic labeling data and additional biochemical approaches, has also shed light onto the mRNA-encoded regulatory mechanisms supporting the synthesis of a hydrophobic protein with multiple transmembrane domains -COX1- and the translation of the bicistronic *ATP8-ATP6* transcript. Moreover, we also studied the mRNA conformational and functional changes induced by LRPPRC, a protein required for mRNA stability, polyadenylation, and translation of most mt-mRNAs. In the absence of LRPPRC, most mRNAs are degraded, but the remaining stable mRNAs adopt different folding patterns compared to wild-type cells. We show that cells lacking LRPPRC have an overall more structured transcriptome, although some regions become more structured while others are more relaxed than in wild-type cells, in a transcript-specific manner. This pattern indicates that LRPPRC may act as a holdase to maintain mRNA folds, which are expected to regulate mt-mRNA stabilization and efficient translation. Altogether, our findings define the mt-mRNA folding landscape, offering insight into post-transcriptional regulation of mitochondrial gene expression. Additionally, we present a strategy that enables the investigation of mt-mRNA folding across cells and tissues and their role in regulating mitochondrial gene expression during development, diseases, and aging.

## Results and Discussion

### Development of mitoDMS-MaPseq to probe mitochondrial genome-wide RNA structures

We aimed to probe the entire mitochondrial mRNA folding landscape in cultured human HEK293T cells using DMS-MaPseq ^28,30^. Our initial attempts using the established method in whole cells failed to provide sufficient read coverage on mitochondrial transcripts to elucidate secondary structures. To address this, we developed and optimized an *in organello* approach. By incubating highly-purified functional mitochondria with DMS followed by a custom rRNA subtraction protocol and a deep-sequencing library preparation ^32^, we achieved a remarkable increase of up to 1000-fold in the reads mapping to mitochondrial transcripts (**Fig. S1A**). The mitoDMS-MaPseq workflow is illustrated in **Figure 1A** and fully detailed in the methods section. Each step required optimization, including DMS concentration, incubation time, and buffer composition to neutralize the extreme acidity of DMS. We found that a DMS concentration of 1% with 5 min incubation was optimal for isolated human mitochondria because longer incubation times could lead to RNA degradation and higher concentrations (*i.e*., 5% DMS) could lead to possible unfolding of RNAs (**Fig. S1B**). In fact, DMS reactivities at 0.5% and 1% DMS correlated very well (R^2^ = 0.93 for *COX1* labeling), while the correlation between labeling with 1% and 5% DMS correlated poorly (R^2^ = 0.56 for *COX1* labeling) (**Fig. S1C**). We also needed to consider that the pH of the mitochondrial matrix is slightly alkaline, close to pH 8, a condition in which DMS probes not only adenines (A) and cytosines (C) but can also probe guanines (G) and uridines (U) ^33^. Because our library preparation method (xGen Broadrange RNA Library Preparation Kit from IDT) is strand-aware, we can separate reads coming from the L and H strands.

**Figure 1.**
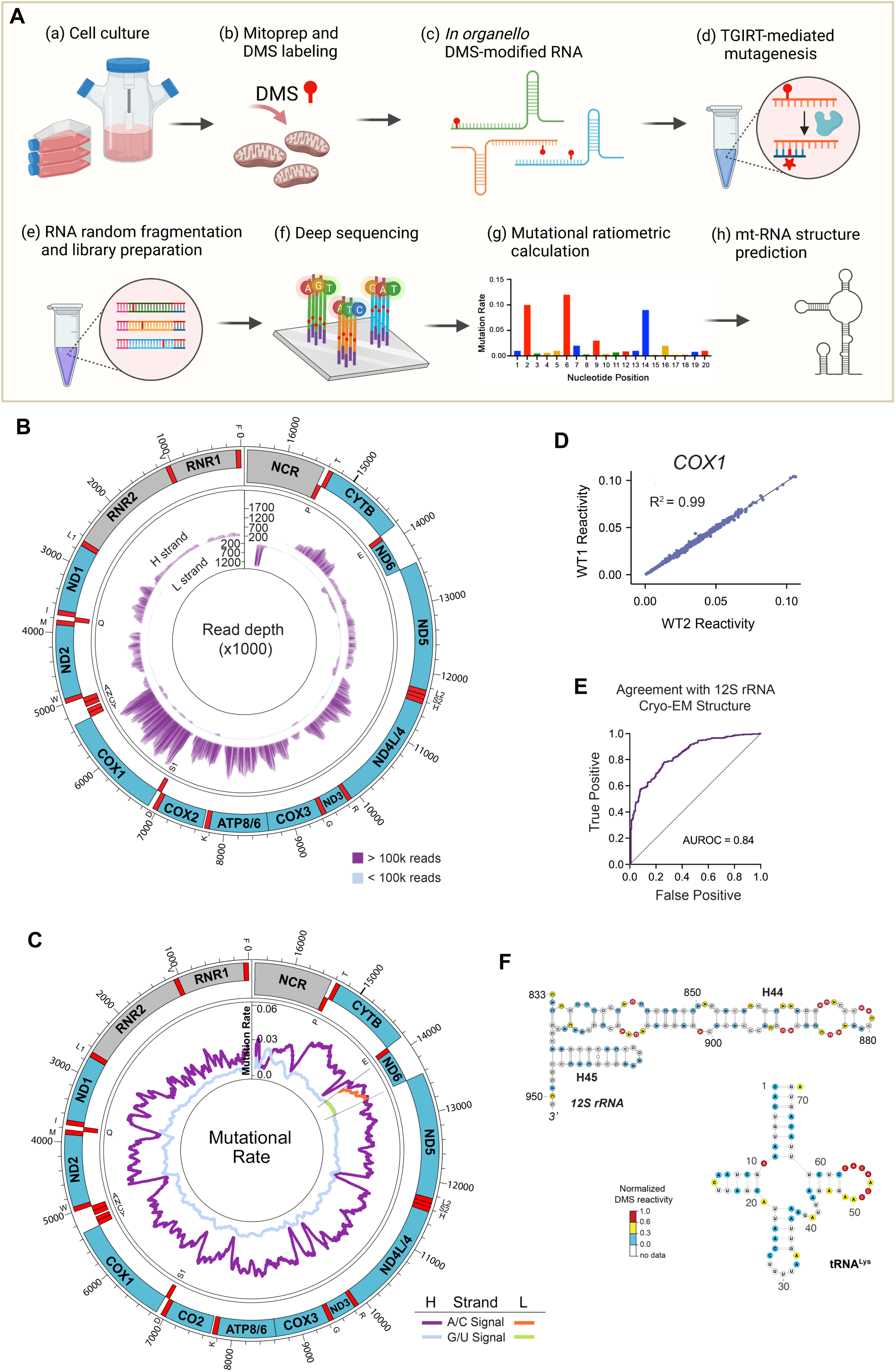
The mitoDMS-MaPseq approach allows for the chemical probing of the mitochondrial mRNA structurome. See also Figure S1. (**A**) Schematic of the mitoDMS-MaPseq workflow. The protocol starts with the DMS modification of RNAs in isolated mitochondria (*in organello*), followed by thermostable group II intron reverse transcriptase (TGIRT)-mediated mutagenesis and library preparation, deep sequencing, and conceptual analysis (see explanation in the text). Created with BioRender.com. (**B**) Circular representation of the mitochondrial genome displaying the number of unfiltered bitvectors across the H-strand and L-strand transcriptomes at each position. (**C**) Circular representation of the mitochondrial genome displaying DMS signal/noise ratio (inner tracks) across the heavy (H) and light (L) strands of the mitochondrial transcriptome. The purple and light blue lines (for the H strand) and orange and green lines (L strand) represent signal *vs*. noise plots of mutation frequencies (*i.e*., among all reads aligning to each strand coordinate, the fraction of reads with a mutation at that coordinate) on adenines (As) and cytosines (Cs) *vs*. guanines (Gs) and uracils (Us) as a function of strand coordinate for untreated and DMS-treated RNA. A mutation frequency of 0.01 at a given position represents 1% of reads having a mismatch or deletion at that position. (**D**) Interexperimental correlation of DMS reactivity for the indicated mitochondrial transcripts at 1% v/v DMS demonstrating high reproducibility. The coefficient of determination R^2^ value is indicated. (**E**) Receiver operator characteristic (ROC) curve comparing the published *12S rRNA* cryo-EM structure ^35^ with DMS reactivities from our datasets. The Area under the curve (AUROC) for DMS reactivity-based prediction is 0.84. (**F**) Pseudo-energy-guided secondary structure model using DMS data on a fragment (H44) of the *12S rRNA* and the *tRNA^Lys^*. Nucleotides are colored by normalized DMS reactivities.

Using the optimized method, we obtained high sequencing coverage across the entire mitochondrial transcriptome (**Fig. 1B**). The samples had high signal-to-noise ratios (**Fig. 1C**), with adenines and cytosines having a mean mutation rate 4.22-fold higher than the background (guanines and uracils) in the mRNAs from both strands (**Fig. 1C**). The DMS reactivity results were highly reproducible between independent biological replicates (*R^2^* > 0.98; **Fig 1D and S1D**). We used the mitoDMS-MaPseq data as constraints in the RNAstructure algorithm ^34^ to fold the entire mitochondrial transcriptome and assessed the quality of our model against two previously-resolved mitochondrial RNA structures, the *12S rRNA* (*RNR1*) ^35^, and mt-*tRNA^Lys^* ^36^ (**Fig. 1E-F**). For that purpose, we first used the area under the receiver operating characteristic curve (AUROC)^29,37^ to evaluate the overall 12S rRNA folding prediction. We determined an AUROC value of 0.84 (**Fig. 1E**), which indicates high-quality probing data considering the complexity and tridimensional folding of rRNAs. We used the *12S rRNA* as an experimental control because although the mitoribosome is highly proteinaceous, the proteins bound to the rRNA do not bind the base side of the RNA nucleotides, allowing for solvent accessibility wherever the RNA is not involved in base-pairing. To further assess the quality of the data, we used the *12S rRNA* H44-H45 region and the *tRNA^Lys^* and determined that the folding prediction matches the known structures (**Fig. 1F**). We thus concluded that mitoDMS-MapSeq reproducibly and accurately probes mitochondrial RNA secondary structures *in organello* on a genome-wide basis.

### mitoDMS-MaPseq provides a model of the population average human mitochondrial mRNA structurome

To analyze mitochondrial transcriptome-wide trends in RNA folding, we used the Gini correlation coefficient, or Gini index of DMS reactivity, which measures the variability in the reactivity of residues as a reflection of RNA secondary structure ^28^. A Gini index close to zero suggests an even distribution of DMS modifications, indicating either unfolded RNA or high structural heterogeneity. Conversely, a Gini index near one implies strong protection of a subset of residues from DMS, indicating a highly stable RNA structure ^31^. In agreement, we measured spikes of Gini index in transcriptome regions known to be rich in secondary structures *in vivo* (rRNAs and tRNAs), which were less frequent and prominent in the coding regions (**Fig 2A**). We used the average mitoDMS-MaPseq data from *in organello* labeling assays as a constraint in RNAStructure^38^ to create a secondary structure model for each mitochondrial mRNA (**Fig. 2B**) and investigated global structurome features.

**Figure 2.**
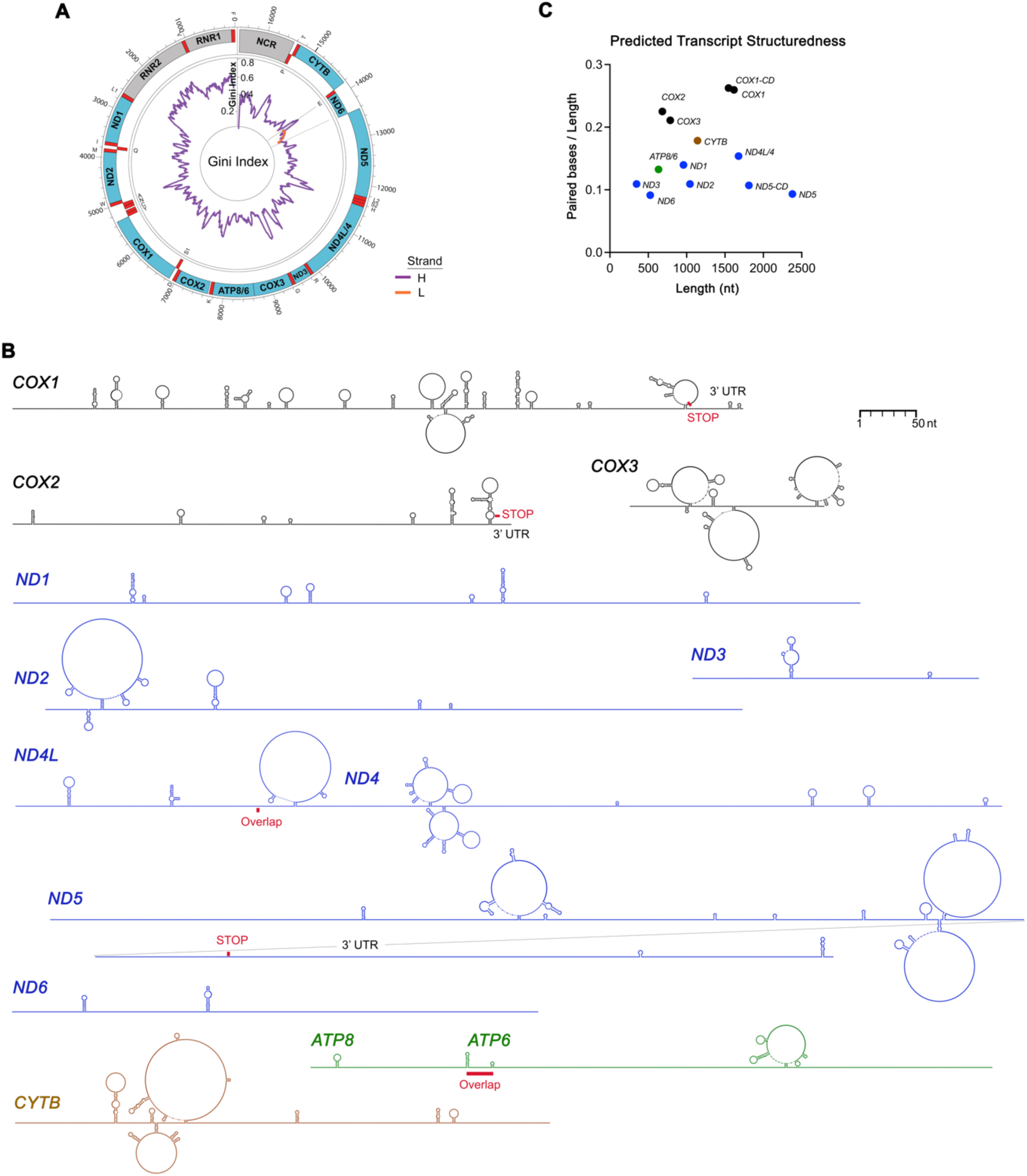
Genome-wide features of the mitochondrial RNA structurome. See also Figures S2 and S3. (**A**) Circular representations of the mitochondrial genome H and L strands displaying the Gini index of DMS reactivity by taking the mean over a sliding window of 80 nt. (**B**) Predicted secondary structure of the 9 monocistronic and 2 bicistronic transcripts that conform the mitochondrial mRNA transcriptome. The 3’ UTR in *COX1*, *COX2,* and *ND5* transcripts are indicated, as well as the overlapping open reading frame regions in the bicistronic transcripts *ND4L/4* and *ATP8/6*. Transcripts are displayed with color-coding corresponding to the OXPHOS complex associated with the encoded protein. (**C**) Predicted transcript structuredness estimated by calculating the number of predicted paired bases normalized by transcript nucleotide length. CD, coding region without the 3’ UTR.

Using the normalized count of predicted paired bases relative to transcript length as a measure of transcript structuredness, we identified *COX1*, *COX2*, and *COX3* as the most structured transcripts (**Fig. 2C**). Intriguingly, these three transcripts were recently reported to undergo the lowest lifetime translation cycles before decay (<10), while the less structured *ND2, ND3, ND5,* and *ATP8/6* transcripts undergo higher lifetime translation cycles (30-60) before decay ^39^. *COX1* and *COX3* exhibit even distribution of stem-loops along the RNA, whereas all other transcripts contain long stretches of linear sequences that characterize the mitochondrial mRNA structurome (**Fig. 2B**). The relatively weak transcript structuredness could be attributed to extensive mitoribosome footprinting during translation. However, mitochondrial Ribo-Seq data calibrated through Ribo-Calibration indicates low mitoribosome occupancy on transcripts, with pools of *ND4L*, *ND6*, *COX3*, and *COX2* mRNAs being mitoribosome-free, and most transcripts carrying only 1-2 mitoribosomes, except for *CYTB* (~3), *ND2* (~4), and *ND5* (~6) ^39^. Hence, the mitoribosome sparseness on transcripts should minimize substantial footprinting effects on the predicted structurome. To assess the impact of the native environment, including factors like RNA-binding proteins, matrix ion composition, ligands, and crowding, on the structure of the mitochondrial mRNAs, we conducted *in vitro* transcription, refolding, and DMS-probing for each transcript to predict their structure. We detected drastic differences in DMS reactivity between transcripts folded *in vitro* and those in native conditions (**Fig. S2A**), underscoring the relevance of our *in organello* approach.

### The 5’ ends of most mitochondrial transcripts are unstructured *in organello*

Unlike prokaryotic mRNA, human mitochondrial mRNAs lack 5’ untranslated regions and Shine-Dalgarno sequences, as well as the typical 5’ methylated cap observed in eukaryotic mRNA, which are all crucial for translation initiation on the small ribosomal subunit. Recent investigations strongly suggest that translation of mitochondrial mRNAs commences on 55S monosomes rather than on the small subunit ^40–42^, akin to the rare leaderless mRNAs in prokaryotes ^43,44^.

Despite these findings, the role of structural elements within the 5’ ends of mitochondrial mRNAs in translation remains uncertain. A sole study that approached this question used *in vitro* transcribed 5’ ends of bovine mitochondrial transcripts (~70 nt in length) and applied SHAPE chemistry ^25^. The study uncovered that while the initial 50 nt of *ND2, ND3*, *COX1*, and *ATP6* were unstructured, the remaining transcripts displayed structures encompassing the initial 20 nt. Nonetheless, these structures were deemed thermodynamically unfavorable, leading to the conclusion that mitochondrial mRNA start codons are situated within structurally accessible, single-stranded sequences that would enhance their recognition by the mitoribosome ^25^.

Our own structural models reveal that in most transcripts, a minimum of ~40 nt downstream of the start codon adopts a linear conformation (**Fig S3**). Three exceptions are *COX2*, *ATP8*, and *ATP6* (**Fig 2B**). *COX2* and *ATP8* are predicted to form stem-loop structures after 16 and 21 nt, respectively, which fall well within the 32-nt footprint of the mitoribosome ^39,45^. These structures, positioned within the crucial region defined by Kozak ^46^, suggest their potential to facilitate translation initiation by impeding scanning, thereby affording additional time for start codon recognition. Notably, *ATP6* start codon (AUG) is in a hairpin that marks the beginning of the *ATP8/6* overlapping region in the bicistronic transcript (**Fig 2B**). This structure could conceivably exert a relevant regulatory role in coordinating *ATP8* translation elongation and *ATP6* initiation at a (−1) frame. Worth noting, the context of the AUG in *ATP6* is unique as the corresponding AUG in *ND4,* within the *ND4L/4* bicistronic transcript, lacks nearby structural elements (**Fig S3**).

### Conformational plasticity of the bicistronic *ATP8/ATP6* transcript supports mechanisms of programmed ribosomal frameshifting (PRF), and Termination-Reinitiation (Te-Re) to ensure the stoichiometric synthesis of ATP8 and ATP6

*ATP6* translation from the *ATP8/6* bicistronic transcript relies on prior translation of the upstream *ATP8* ORF, but not *vice versa* ^47^. The ribosome cannot load directly onto the *ATP6* −1 ORF, probably due to constraints imposed by the long 5’ UTR ^25^. This reveals a unidirectional dependence, coupling the translation of the two ORFs. Notably, despite ATP6 and ATP8 being subunits of the F_1_-F_o_-ATP synthase and existing in a 1:1 ratio, mitoribosome profiling studies have indicated double the mitoribosome occupancy for *ATP8* compared to *ATP6* in the transcript ^45^, along with a double number of translation cycles before decay ^39^. These findings imply the presence of translational or post-translational mechanisms to uphold the 1:1 stoichiometry.

mitoDMS-MaPseq and DREEM clustering analysis unveiled two distinct conformations within the overlapping region (46 nucleotides long) of the *ATP8/6* bicistron (**Fig 3A**). Among the mRNA population, 35% exhibited an unstructured region, while the remaining 65% displayed a stem-loop structure obstructing the *ATP6* start codon. The hairpin is preceded by a heptanucleotide sequence 5’-AAAAAAU-3’ and a 23-nt spacer sequence within the *ATP8* ORF (**Fig 3B**). These structural features strongly suggest the existence of a programmed ribosomal frameshifting (PRF) mechanism, likely governing *ATP8* and *ATP6* translation. PRF is a translational recoding mechanism prevalent in organisms with compact genomes such as viruses ^48^ and rare in bacteria or eukaryotic cells ^49^. In PRF, the translating ribosome slips into an alternative open reading frame (ORF) during translation in a regulated manner ^50^, enabling the translation of multiple proteins from a single transcript. PRF usually arises from the effects of two cis-elements within the transcript, as seen in the *ATP8/6* bicistronic transcript: a ribosomal “slippery” heptanucleotide sequence at which −1 PRF occurs, followed by a downstream stimulatory RNA structure, such as a stem-loop or pseudoknot. When encountering this block, the translating ribosome pauses, allowing for slippage and subsequent PRF. A PRF mechanism on the *ATP8/6* transcript could explain the unidirectional translation dependency ^47^ and differential ribosome occupancy in the two ORFs ^45^.

**Figure 3.**
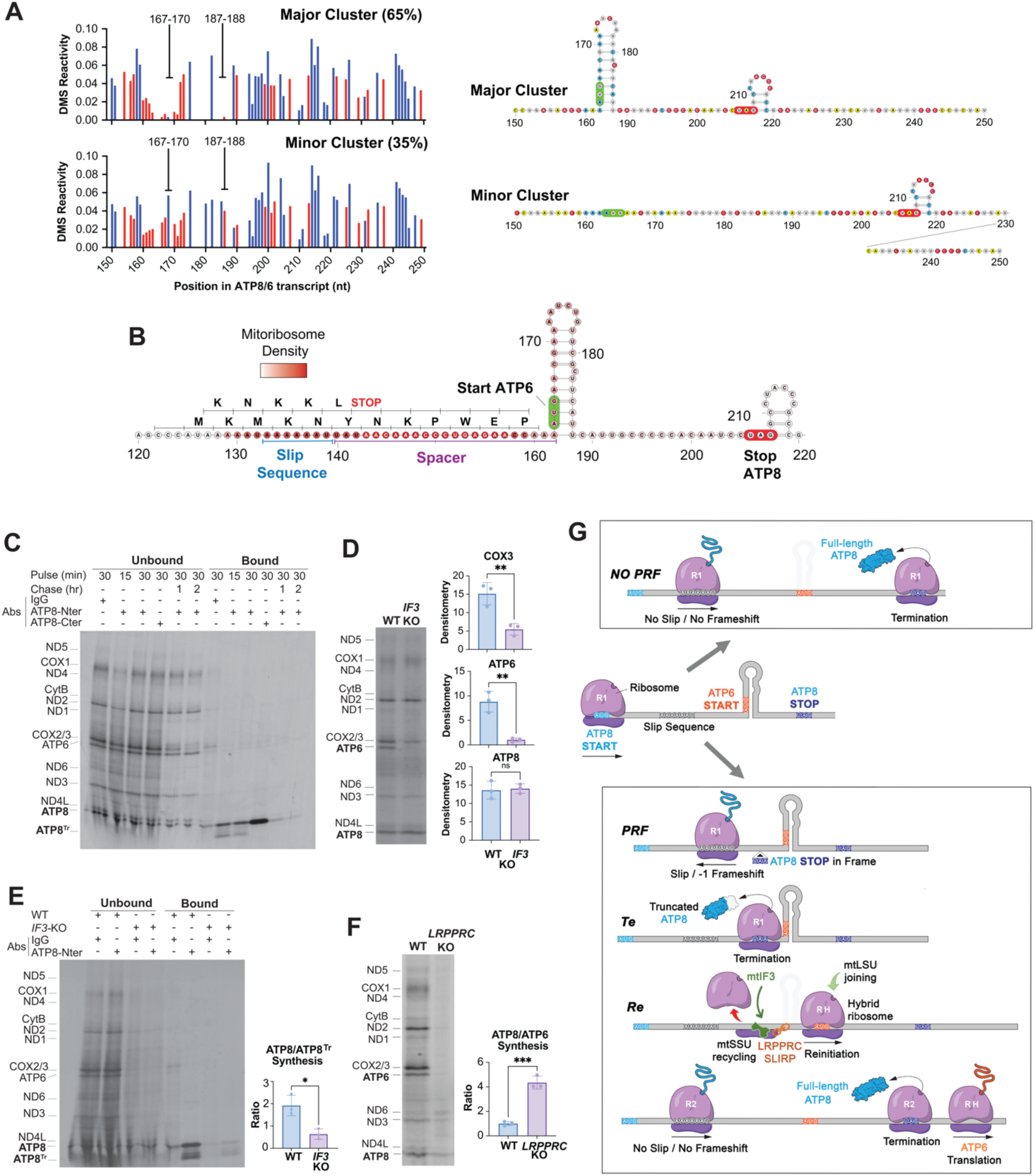
Programmed ribosome frameshifting and termination-reinitiation govern the translation of the two ORFs encompassed in the ATP8/ATP6 bicistronic transcript. See also Figure S4. (**A**) DREEM-driven clustering analysis of the Mito-DMSMaPseq data on the *ATP8/6* bicistronic transcript. For each cluster, the DMS-reactivity plot is shown on the left panel, and the predicted secondary structure is shown on the right panel. (**B**) Predicted secondary structure of the *ATP8/6* mRNA overlapping region containing the hairpin that occludes the start codon of *ATP6* (highlighted in green). The red coloring indicates overlapped published ribosome profiling data ^9^. The location of the potential slippery and spacer sequences is marked. (**C**) Metabolic labeling of mitochondrial translation products with [S^35^]-methionine in mitochondria isolated from WT cells (*in organello*) followed by immunoprecipitation with antibodies against either the N- or the C-terminus of ATP8, or IgG as a control. In the indicated lines, a chase period was included. (**D**) Metabolic labeling of mitochondrial translation products with [S^35^]-methionine in WT and *IF3*-KO cells (*in cellula*) in the presence of emetine to inhibit cytosolic protein synthesis. The graphs show the quantification of COX3, ATP6, and ATP8 in 3 independent assays. T-test, **p<0.001. (**E**) *In organello* protein synthesis and immunoprecipitation assay in WT and *IF3*-KO cells, performed as in panel (C). The graph shows ATP8/ATP8^Tr^ ratio quantification in 3 independent assays. T-test, **p<0.001. (**F**) *In cellula* mitochondrial protein synthesis in WT and *LRPPRC*-KO cells. The graph shows the ATP8/ATP6 ratio quantification in three assays. T-test, **p<0.001. (**G**) Model depicting the translation mechanism of ATP8 and ATP6 from the bicistronic transcript.

The co-existence of two *ATP8/6* folding variants highlights conformational plasticity, potentially linked to the translation mechanism. In both variants, the mitoribosome will load onto the 5’ of the *ATP8* ORF and initiate translation. In the transcript lacking the hairpin enclosing the *ATP6* AUG, the mitoribosome would traverse the slippery sequence without finding a downstream block, resulting in exclusive ATP8 synthesis. Conversely, in the structured transcript, when encountering the hairpin, the mitoribosome pauses and walks, as evident in the prominent mitoribosome profiling signature, found in published datasets ^9,45^, in the *ATP8/6* overlapping region and adjacent upstream sequence (**Fig 3B**). Following the pause, two possibilities arise. The mitoribosome could go through the loop and terminate ATP8 synthesis. Alternatively, the pause might induce mitoribosomal slippage and −1 PRF. However, as the mitoribosome continues to read through after −1 PRF, it would encounter either a UAA or AGA stop codon before reaching the *ATP6* start codon, producing a truncated ATP8 polypeptide, lacking the C-terminal 21 amino acids (**Fig 3C**). In agreement with this prediction, using *in organello* protein synthesis followed by immunoprecipitation with antibodies against ATP8, either its N-terminal or C-terminal ends, we detected the truncated ATP8 polypeptide (ATP8^Tr^) solely with the anti-N-term antibody and established that it is fast degraded in chase experiments (**Fig 3C**).

After synthesizing the truncated ATP8, a termination-reinitiation (Te-Re) mechanism ^51,52^ must facilitate *ATP6* ORF translation. Following premature *ATP8* ORF termination, the mitoribosome requires a means to remain tethered to the mRNA, scanning for a nearby start codon until encountering the *ATP6* AUG (**Fig 3B**). Te-Re is notably common in bicistronic viral protein translation. This process can entail both cis and trans-acting elements that preserve either the small subunit (mtSSU) or the entire post-termination monosome associated with the mRNA, enabling forward scanning for the next ORF. In mitochondria, translation of *ATP6* ORF specifically requires the translation initiation factor 3 (mt-IF3) ^42,53^ (**Fig 3D**), and its absence lowers the ATP8 *vs* ATP8^Tr^ proportion by ~3-fold (**Fig 3E**). The dependency of *ATP6* translation on mt-IF3 points toward mtSSU recycling. Unlike caliciviruses, which use a sequence element denoted ‘termination upstream ribosomal binding site’ (TURBS) to retain the SSU via interaction with helix 26 of the 18S rRNA ^54^, *ATP6* reinitiation employs a distinct mechanism as these RNA features lack mitochondrial counterparts. A candidate tether of the mt-SSU with the *ATP6* 5’ UTR is the LRPPRC-SLIRP complex, known to interact with mRNAs on the mtSSU ^24^ and bind the *ATP8* ORF just at the 5’ of the putative slippery sequence ^23^. Supporting this notion, the ATP6/ATP8 synthesis ratio drops significantly by a factor of four in an *LRPPRC*-KO cell line (**Fig. 3F**), despite 79% of *ATP8/6* transcripts exhibiting the hairpin in the ORFs overlapping region (**Fig. S4A**).

The two *ATP8/6* transcript conformations identified *in organello* may be interconnected, as only the structured conformation arises *in vitro* (**SD10**). We propose a model (**Fig. 3G**) in which a translating ribosome, as it resolves and traverses the RNA stem-loop structure, unfolds the transcript. A subsequent trailing ribosome utilizes this unfolded state to synthesize ATP8 before the structure reforms. This concept is supported by the observation that a spike of mito-disomes was detected in the bicistronic transcript with the leading mitoribosome positioned at nt 195 in the overlapping *ATP8/6* region and the trailing mitoribosome within the *ATP8* ORF ^39^. The relative load of ribosomes on the transcript affects the propensity of ribosomes to collide, which is expected to influence the efficiency of PRF in the stalling and trailing ribosomes, akin to viral systems ^55^.

Notably, the primary sequence surrounding the ATP6 starting site has a high degree of conservation among animal mtDNAs (**Fig. S4B**), suggesting that the PF mechanism could be conserved in these species.

### *The COX1* secondary structure supports mRNA-programmed translational pausing

In all living organisms, ribosomes translating membrane proteins are targeted to membrane translocases early in translation. Mechanisms exist to modulate protein folding by regulating the translation elongation rate, including mRNA sequence elements as well as cis- and trans-acting factors that influence ribosome speed ^56–59^. The translating ribosome must unfold mRNA secondary structures to feed single-stranded mRNA through the narrow mRNA channel. The bacterial and eukaryotic ribosomes are efficient helicases ^60^, explaining why most secondary structure elements within coding regions of mRNAs do not influence the translation elongation rate ^61^. However, specific mRNA stem-loop structures can induce ribosome pausing or stalling through a mechanism known as mRNA-programmed translational pausing, which significantly impacts the synthesis of hydrophobic membrane proteins ^62,63^. Notably, ribosome profiling studies have shown that in many bacterial and eukaryotic hydrophobic proteins with multiple transmembrane domains (TMs), ribosomes prominently pause before TM2 once TM1 has emerged from the ribosome and is inserted into the membrane (*9*). In human mitochondria, membrane-bound mitoribosomes recruit the OXA1L translocase machinery during protein synthesis initiation to facilitate the co-translational insertion of hydrophobic proteins with multiple TMs ^64^. The likelihood of mRNA-programmed translational pausing might be heightened by the absence of certain proteins (full uS4, specific domains of uS3, and bS1) in the human mitoribosome, which in bacteria aid in mRNA unfolding during translation. Although mitochondria have evolved a system for mRNA delivery based on transacting proteins (LRPPRC-SLIRP) and mS39 acting as a linker, these are PPR motif-containing proteins without known helicase activity ^24^.

We have examined the correlation between mRNA hairpin distribution and the location of TM coding sequences. Although no universal rule applies to all transcripts and TMs, certain patterns have emerged. For instance, in *COX3*, *CYTB*, and *COX1* (**Fig. 4** and **S5**), several TM-coding regions are flanked by hairpins. Mito-disome accumulation during the translation of some of these sequences ^39^ suggests the occurrence of mRNA programmed translational pausing. In *ND6*, mito-disomes accumulate after the synthesis of TM1 and TM3, corresponding to the positions of the two single mRNA hairpins in the transcript (**Fig. S5A**). A similar correlation between protein structure and neighboring mRNA structure has been previously reported in several organisms and suggested to occur because structured RNA regions slow ribosome processivity and thus facilitate correct folding of the preceding TM protein domains ^65–67^. In mitochondria, the lack of full correlation between transcript and protein structures suggests that when it occurs, it may be for specific regulatory purposes that remain to be unveiled.

**Figure 4.**
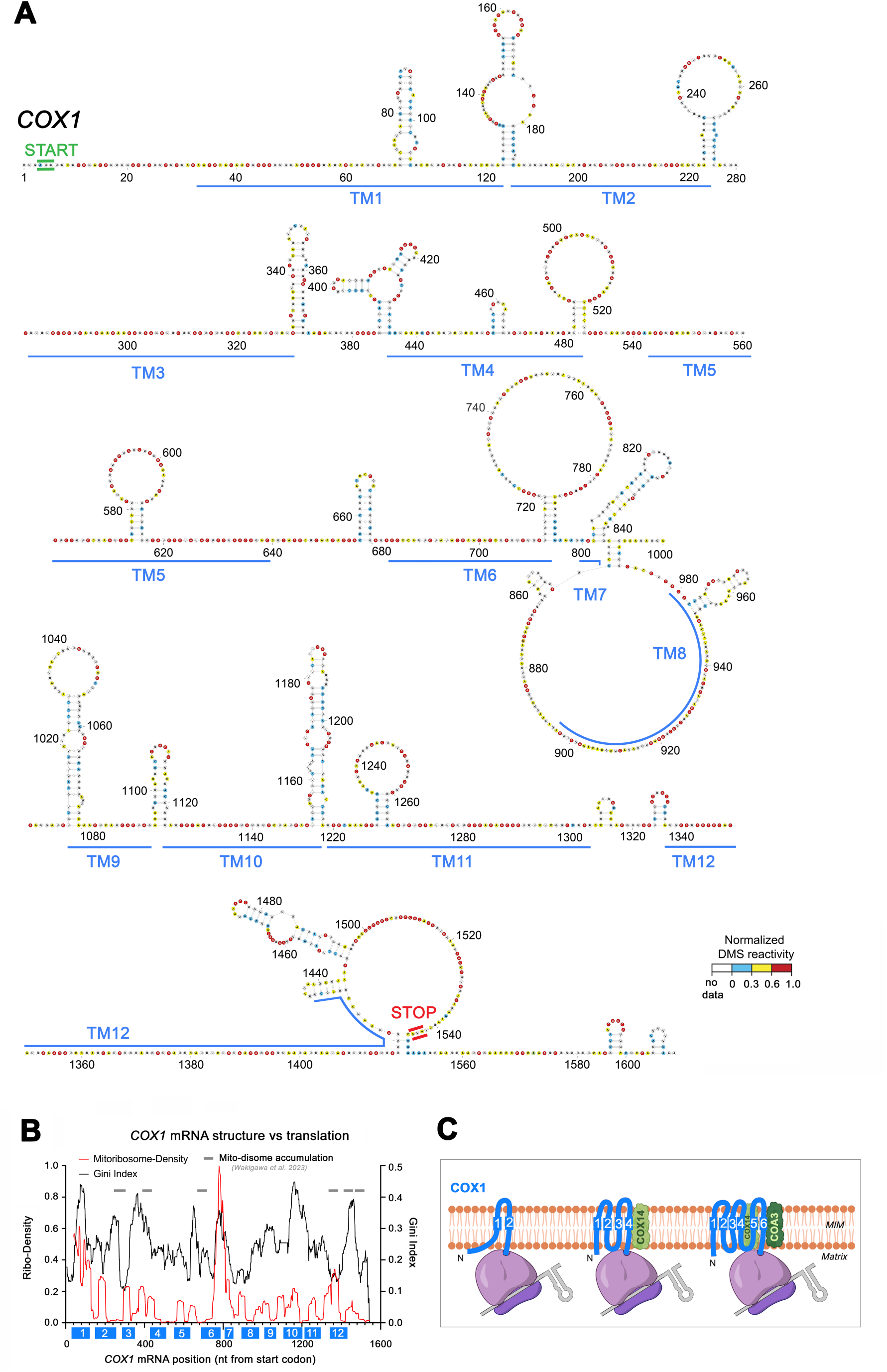
Secondary structures in the *COX1* transcript flank regions coding for transmembrane domains. See also Figure S5. (**A**) Predicted secondary structure of the *COX1* mRNA. Nucleotides are colored by normalized average population DMS reactivity. The regions coding for transmembrane (TM) domains in the COX1 protein ^77^ are indicated with blue lines. (**B**) Analysis of the *COX1* mRNA structure vs. translation by plotting the Gini correlation index (structure) and mitoribosome density profiles (retrieved from published ribosome profiling data ^9^) across the *COX1* mRNA sequence. Regions where mito-disomes have been reported ^39^ are marked with grey lines. The sequences coding for COX1 TMs are indicated as blue blocks. (**C**) Schematic showing co-translational insertion of COX1 TM domains and how secondary structure in the mRNA allows for co-translational pausing to allow for TM domain insertion. Previously reported binding of COX1 chaperones COX14 and COA3 to newly synthesized COX1^68^ is depicted.

We focused on COX1, encompassing 12 TM helices, and recognized for its well-studied TM biogenesis. The secondary structure of *COX1* mRNA has revealed the presence of stem-loops in most segments encoding TM helices, either containing or flanking them (**Fig 4A**), capable of inducing translational pausing (**Fig 4A**). Translation of *COX1* mRNA is linked to the import of nucleus-encoded cytochrome *c* oxidase subunits to facilitate co-translational folding, stabilization, and assembly of COX1 through a translational plasticity pathway ^68^. During synthesis, the assembly factors COX14 (C12ORF62) and COA3 (MITRAC12) sequentially associate with the nascent COX1 chain ^68^. Previously, three major puromycin-released *COX1* mRNA translation truncated products were detected having 2, 4, or 6 TMs (^68^ and **Fig. 4B**). The pauses after TM4 or TM6 have been attributed to COX14 and COA3 binding, respectively ^68^. In agreement, published mitoribosome profiling data detected pauses at TM2, TM4, and all other TMs, most prominently at TM6-7 ^9,39^ (**Fig. 4B**), and mito-disomes accumulate post-synthesis of TM2 and prior to TM4 synthesis ^39^ (**Fig. 4A-B**). The *COX1* mRNA region spanning domains encoding TM6 and TM11 is notably structured (**Fig. 4A**). This region also involves frequent mitoribosomes collisions ^39^, probably to pace elongation rate to membrane insertion and folding of the COX14-COA3-bound nascent COX1 polypeptide. We conclude that COX1 synthesis follows a discontinuous pattern facilitated by *COX1* mRNA-programmed translational pausing. This mechanism underpins a translational plasticity system that allows COX1 protein synthesis to adapt to the influx and assembly of nucleus-encoded subunits ^68^.

### Stop codons of *COX1*, *COX2* and *COX3* are positioned within structured domains

The three mRNAs coding for cytochrome *c* oxidase subunits have their stop codon lying near or within structural features (**Fig. 2B**, **5A**, and **S5**). COX2 and COX3 have canonical UAA and UAG stop codons respectively, which are recognized by the release factor mtRF1a. Instead, *COX1* has a non-canonical AGA codon that is decoded by mtRF1 ^69–71^. The specific recognition of the noncanonical stop codon is complex ^71^ and probably is a slow process, given the queue of mitoribosomes forming before and at the stop codon ^39^. Notably, preceding the stop codon, two significant stem-loops exist (**Fig. 4A**), potentially contributing to the queuing phenomenon. *COX1* is one of the three mRNAs having a 3’ trailer (72 nt), which concludes with two small hairpins (**Fig. 4A**). These structures may help to promote the reported accumulation in this region of mito-disomes, likely containing unrecycled mitoribosomes pushed by translating trailing units ^39^. Queuing and mito-disome accumulation does not occur in *COX2*, but it does just before the *COX3* stop codon ^39^, although in this case the region codes for the last TM of the COX3 protein (**Fig. S5A**), which may influence the pausing.

**Figure 5.**
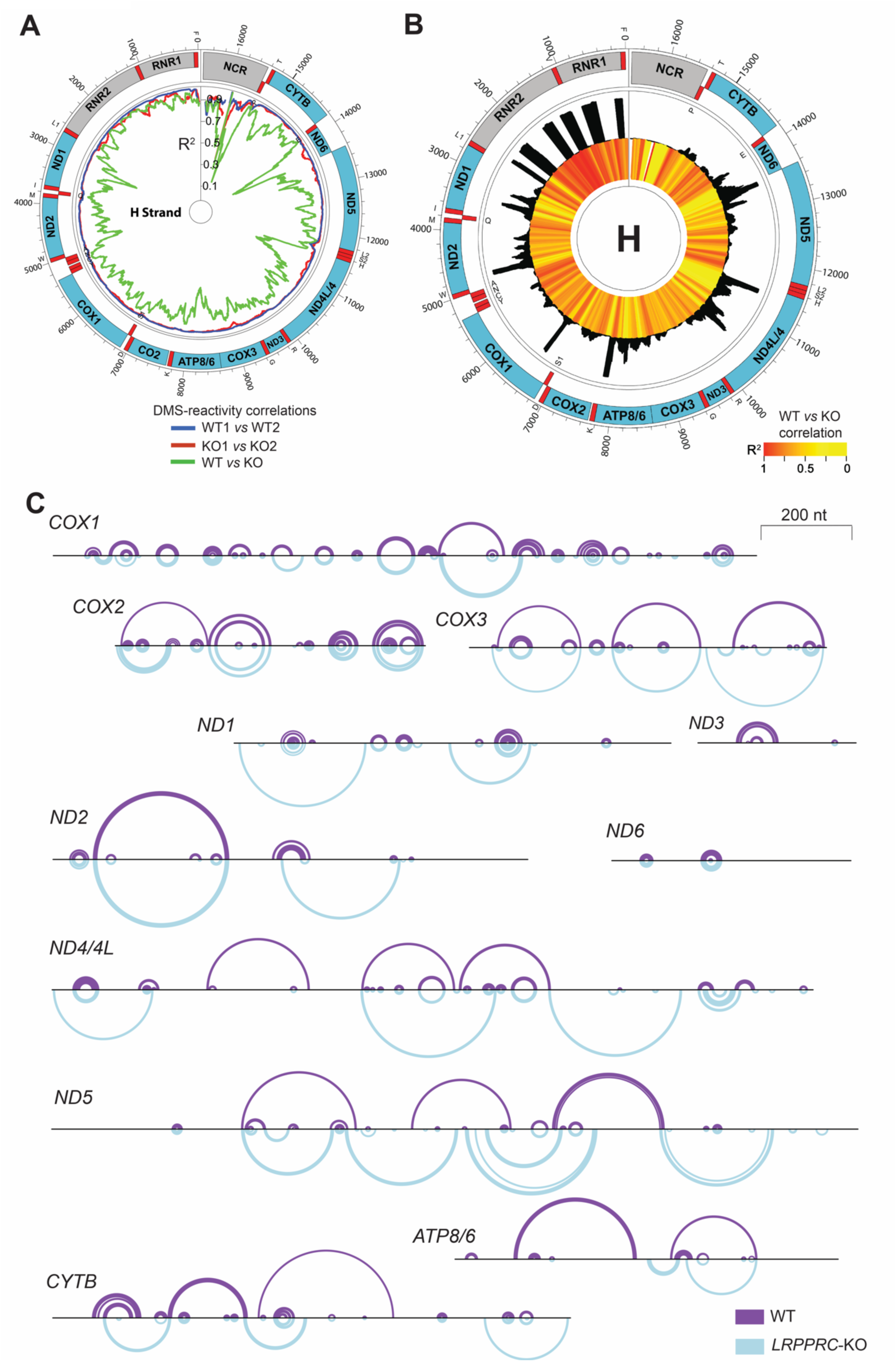
A remodeled mitochondrial mRNA structurome arises in the absence of LRPPRC. See also Figures S6 and S7 and supplemental datafiles SD1-SD10. (**A**) Circular representation of the mitochondrial genome H-strand displaying the indicated DMS-reactivity correlations between the WT and *LRPPRC*-KO structuromes. (**B**) Circular representation of the mitochondrial genome H-strand displaying published LRPPRC binding sites as determined by PAR-CLIP ^23^ (black histograms) and changes in DMS reactivity in the *LRPPRC*-KO cell lines as compared to WT (red-yellow shades reflecting R^2^ values). (**C**) Linear arc plots indicating predicted areas of secondary structure changes within each mitochondrial transcript between WT and *LRPPRC*-KO mitochondria.

### A divergent mt-mRNA structurome emerges in the absence of LRPPRC

The mitoDMS-MaPseq method enables the exploration of interactions between mt-mRNAs and proteins, alongside associated conformational and functional shifts in transcripts. This is exemplified by LRPPRC, a protein implicated in the French-Canadian variant of Leigh’s syndrome —a childhood neurodegenerative disorder linked to mitochondrial respiratory chain CIV deficiency ^22^. LRPPRC ensures stability, polyadenylation, and translation of all mt-mRNAs except ND6 ^23,72^. Previous RNAseq and mitoribosome profiles indicated distinct functions of LRPPRC in mRNA stability and mitoribosome targeting ^45^. By comparing WT and *LRPPRC*-KO structurome datasets, we have now investigated potential links between mRNA structural remodeling, stability, and translation efficiency.

Using mitoDMS-MaPseq in *LRPPRC*-KO mitochondria, we obtained high sequencing coverage across the entire mitochondrial transcriptome (**Fig. S6A**). The samples had high signal-to-noise ratios in the mRNAs from both strands (**Fig. S6B)**. As for the WT data, the DMS reactivity results were highly reproducible between independent *LRPPRC*-KO biological replicates (*R^2^* > 0.95; **Fig. S6D**), and the Gini index of DMS reactivity displayed spikes in transcriptome regions corresponding to rRNAs and tRNAs, which were more scarce and less prominent in the coding regions (**Fig S6C**).

Importantly, when correlating population average DMS reactivities between the WT and KO in each individual mitochondrially encoded transcript, we detected a poor correlation between the WT and *LRPPRC*-KO data for all mRNAs (*R^2^* = 0.47-0.73). Remarkably, ND6 stood out as the sole exception, maintaining a consistent DMS signal (*R^2^* = 0.93). This result is in agreement with previous literature showing that *ND6* is the single mt-mRNA not bound by LRPPRC ^23,72^ (**Fig. 5A and S6E**). Enrichment of differential mitoDMS-MaPseq counts across transcripts (**Fig. S7A**), and comparison of the WT and *LRPPRC*-KO population average models using the mean of sensitivity and positive predictive value (PPV), the Fowlkes-Mallows index (FMI) ^73^, confirmed the differences between the two transcriptomes except for *ND6* (**Fig. S7B**). Although the total number of predicted paired bases was not markedly different across the WT and *LRPPRC*-KO transcriptomes for most mRNAs, the number for *LRPPRC*-KO was lower in *COX1*, *ATP8/6*, and *ND3* transcripts and higher in *ND5*.

Notably, overlaying the DMS-reactivity differences between the WT and the *LRPPRC*-KO contexts onto established LRPPRC binding sites ^23^ revealed a noteworthy connection (**Fig. 5B**), suggesting the intriguing possibility of having detected LRPPRC-induced RNA footprinting in the WT structurome. This would imply that the binding affinity of LRPPRC lies with the base of the RNA molecule rather than its sugar backbone, which is not common among conventional RNA binding proteins. However, cryo-EM analyses of the LRPPRC/SLIRP-mRNA complex bound to a translating mitoribosome, showed interaction with the backbone of the few resolved nucleotides ^24^. Moreover, *in organello* DMS-reactivities in *LRPPRC*-KO markedly differ from those of *in vitro* folded mRNAs (*R^2^*=0.09-0.51 for individual transcripts) (**Fig. S2B**), which was also broadly reflected in the number of predicted paired bases and the FMI for most transcripts (**Fig. S7B-C**). Importantly, if the binding of LRPPRC reduced DMS accessibility, we would measure that as a global increase in structure. Instead, many bases are more accessible to DMS in the WT compared to the *LRPPRC*-KO, and we observe a change in structure that is supported by thermodynamic models (**Fig. 5C** and **S7D**). These observations minimize the potential reflection of LRPPRC footprinting in the WT structurome.

A previous study using isolated mouse heart mitochondria subjected to endogenous single strand specific RNAse A and Rnase T1 footprinting, and Rnase If footprinting analyses, reported that loss of LRPPRC results in a net increased protection against cleavage by these endonucleases suggesting LRPPRC generally prevents folding of mitochondrial mRNAs ^23^. Our models of transcript structures in the *LRPPRC*-KO also indicate deviations from WT structures occurring across transcripts (except *ND6*) but including both loss and formation of new stem-loops (**Fig. 5C** and **S7D**). A side-by-side comparison of the predicted secondary structure of each transcript in WT and *LRPPRC*-KO mitochondria and *in vitro* folded mRNAs is presented in the supplementary datafiles **SD1-SD11**. Therefore, we propose that LRPPRC does not necessarily act to prevent secondary structure formation where it binds ^23^, but rather remodels mRNA folding presumably to facilitate its stability and translation.

### DREEM-based clustering analyses reveal an ensemble of secondary conformations across the mitochondrial mRNA transcriptome

RNA molecules often exist in a dynamic equilibrium, comprising an ensemble of alternative structural forms ^37,74,75^. The conformation of each RNA is dictated by a thermodynamic landscape that changes with the RNA sequence and the environmental conditions. Depending on these factors, a given RNA populates an ensemble of conformations of various compactness, stabilities, and flexibilities ^75,76^. The functional relevance of the alternative secondary structures we have described for the *ATP8/ATP6* overlapping region (**Fig. 3G**) underlines the importance of defining potential conformational ensembles across the mitochondrial transcriptome. For *ATP8/6*, the average DMS signal supported a hairpin structure in that region in WT but not in *LRPPRC-KO* mitochondria (**Fig. SD10**). However, through clustering deconvolution, it became evident that the hairpin structure is indeed prevalent in 65% and 79% of the respective mRNA molecules.

For simplicity, all structures presented and discussed so far, except for the *ATP8/6* transcript, are based on DMS reactivity profiles that reflect an averaged measurement encompassing all potentially co-existing conformations for each given mitochondrial transcript. To assess the accuracy of the predicted structures based on average population DMS reactivity data, we employed AUROC, as previously documented ^37^. It is important to note that a low AUROC (<0.8) signifies a significant disagreement with the structure model, while a high AUROC supports but does not necessarily confirm the correctness of a predicted structure ^37^. Consequently, AUROC served as a metric for identifying structures that lacked strong support from the underlying population average data, implying the potential presence of alternative conformations. The mean AUROC values proved to be robust for both the WT (0.918) and *LRPPRC*-KO (0.913) mRNA transcriptomes (**Fig. 6B** and **6D**), indicating an overall excellent structure prediction performance. Across the mRNA transcriptome, approximately 35% and 49% exhibited AUROC values below the mean in the WT and *LRPPRC*-KO cell lines, respectively, suggesting the existence of multiple potential structural arrangements. From these regions, 15% in the WT and 21% in the *LRPPRC*-KO had AUROC lower than 0.85, and 8% and 12%, respectively, lower than 0.8, which could be the best candidates to undergo alternative folding.

**Figure 6.**
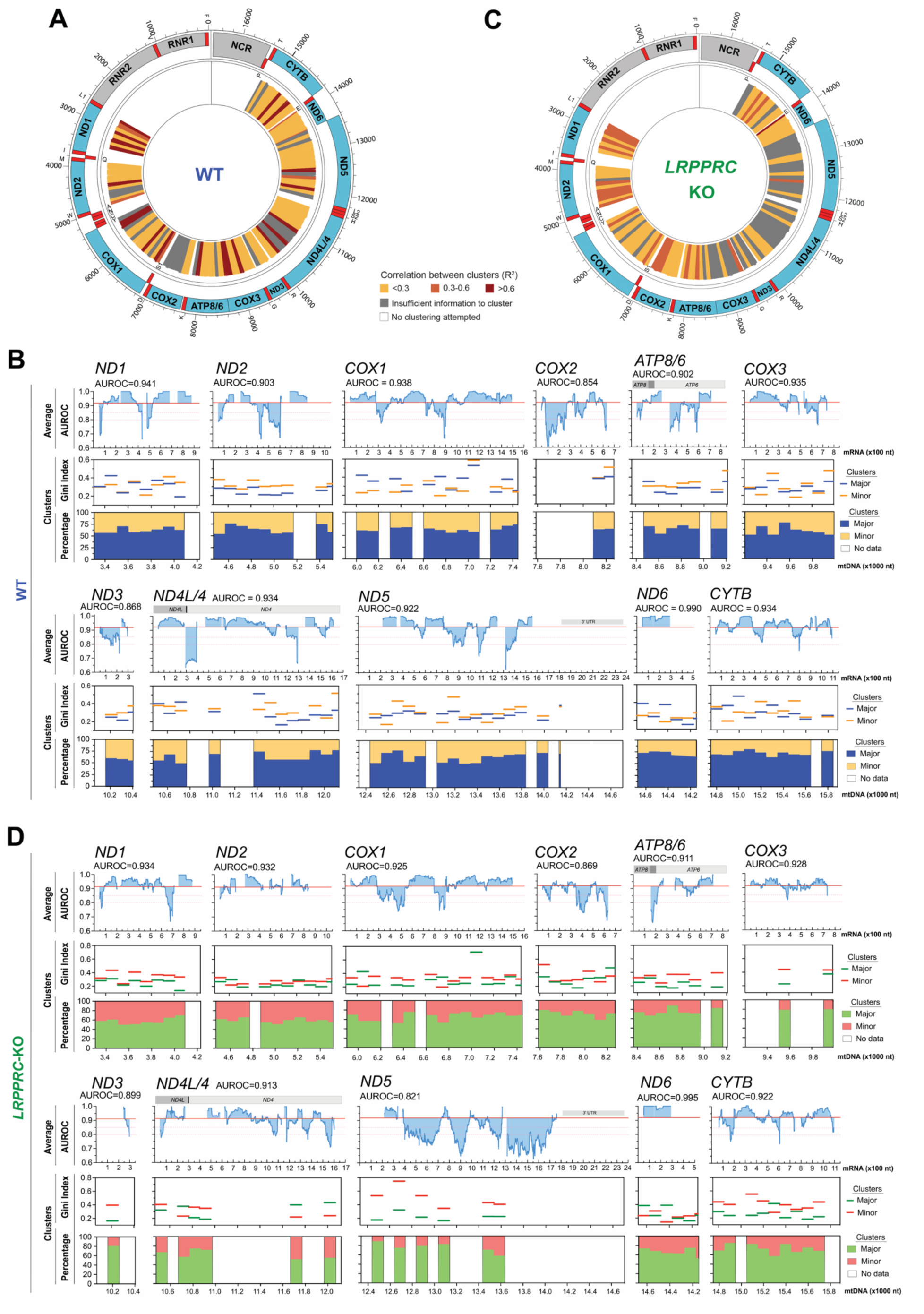
An ensemble of secondary structures populates the mitochondrial transcriptome. (**A** and **C**) Circular representation of the mitochondrial genome displaying the DREEM clustering analysis of DMS-reactivities across the WT (**A**) and *LRPPRC*-KO (**B**) transcriptomes. The regions with enough reads for clustering analysis and the correlation between the two clusters determined in each informative 100 nt window are presented. (**B** and **D**) Linear representation of each transcript showing the agreement between DMS reactivities and predicted secondary structures (AUROC, blue) for the mt-mRNA transcriptome-wide models in WT and *LRPPRC*-KO mitochondria. AUROC was calculated over sliding windows of 100 nt in 1-nt increments; *x* values represent the centers of the windows. The area between the local value and the transcript median is shaded. Regions with AUROC below 0.8, indicating suboptimal modeling, are indicated with pink lines. The two lower panels represent clustering parameters, the percentage of the major and minor clusters (bottom graph), and the Gini index of DMS reactivity (middle graph) for each cluster across 100 nt non-overlapping sliding windows.

To investigate alternative conformations of mt-mRNAs, we applied the DREEM algorithm ^31^ to our *in-organello* mitoDMS-MaPseq dataset using 100-nt non-overlapping sliding windows across the transcriptome and with the algorithm set to look for two clusters. Approximately 75% and 80% mRNA regions were covered by enough sequencing reads (>10^5^ reads) to allow clustering in WT and *LRPPRC*-KO mitochondria, respectively (**Fig 1B, 6A** and **6C**). With this strategy, we covered most regions with low AUROC. Within all regions, the major cluster contained, on average, ~60% of the reads (range between 55 and 75% for the WT and 55-85% for the *LRPPRC*-KO), and the data defining the two clusters for each window with low AUROC exhibited in most cases, low correlation (**Fig 6A** and **C**), which was reflected in their disparate Gini indexes (**Fig 6B** and **D**). These clustering profile parameters underscore the potential prevalence of alternative structures. For each transcript, the two clusters identified in every window were categorized as major or minor based on their percentage contribution to the total. However, it is crucial to emphasize that this categorization does not imply the existence of merely two alternative structures for each transcript; rather, it suggests the potential ensemble of combinations among the clusters within each window. In all examined transcripts, we detected regions where clusters displayed markedly distinct Gini indexes, indicating large differences between these clusters (**Fig 6B** and **D**). We noticed that in WT mitochondria, approximately 23% of the windows had major clusters with Gini indexes at least 0.15 higher than those in the minor clusters, while roughly 33% displayed the opposite pattern. In contrast, in *LRPPRC*-KO mitochondria, only around 15% of the windows displayed major clusters with Gini indexes at least 0.15 higher than those in the minor clusters, while roughly 67% displayed the reverse trend. These data indicate that LRPPRC tends to preferentially stabilize or capture regional mRNA conformations from the ensemble with higher Gini Indexes or “more structured”.

## Conclusions

Mammalian mitochondria have evolved sophisticated mechanisms to regulate gene expression, from transcription to mRNA stability, modification, translation, and degradation. Translation occurs in dedicated mitoribosomes, which are the center of a machinery with features that determine translation control, such as leaderless mRNAs, bicistronic transcripts, or mRNA-binding proteins that hand off the mRNAs to the mitoribosome. Our work provides the first insight into the ensemble of secondary structures of an entire mitochondrial transcriptome. The human mt-mRNA folding landscape we have described constitutes a new layer of biological information. Furthermore, we illustrate how LRPPRC, a broad mt-mRNA protein-ligand that participates in multiple mt-mRNA transactions, shapes this structurome. Finally, the work discloses several critical examples of how mRNA structure guides translation and paves the way toward a complete understanding of the mtDNA-wide and transcript-specific relationships between mRNA structure and its stability, processing, or translation across cell lines, and changing metabolic and environmental conditions.

## Limitations of the study

Our study was performed *in organello* using mitochondria maintained translationally competent, isolated from cultured HEK293T cells. Despite the quality of our mitochondrial preparations, there is always a chance that the purification procedure could perturb mRNA structures. Although the bulk of the structures we have resolved are expected to be conserved across cell lines, there can be some differences in the mitochondrial mRNA landscape depending on how reliant the cells or tissues are on aerobic metabolism and the rate of mitochondrial gene expression. Differences could also be associated to mtDNA haplotypes, which are specific combinations of genetic variations or single nucleotide polymorphisms (SNPs) found within the genome that represent large branches of the human mitochondrial phylogenetic tree. The secondary structures populated in the mRNA folding ensembles that we have described are expected to be influenced by intracellular and environmental conditions, and, therefore, may change across cell lines and tissues. Our study informs about mRNA secondary structures, and all modeling highlights the importance of Watson–Crick base pairing. However, caution is required when using Watson–Crick pairing synonymously with “RNA structure”, given the additional factors influencing natively folded RNA three-dimensional structures. Another relevant limitation relates to our data analysis parameters, which were not set up to detect long-range interactions that usually span distances between a few hundred and several thousand nucleotides. Suspected long-range interactions could be addressed by using antisense oligonucleotides, but their import into mitochondria is challenging. A recent study has used morpholinos to disrupt mRNA-specific protein synthesis *in organello* ^47^, an approach we plan to implement in the future. This approach will also be useful to provide further proof for the programmed translational frameshifting and termination-reinitiation models that we are proposing.

## Supporting information

Supplemental Datafile 1

Supplemental Datafile 2

Supplemental Datafile 3

Supplemental Datafile 4

Supplemental Datafile 5

Supplemental Datafile 6

Supplemental Datafile 7

Supplemental Datafile 8

Supplemental Datafile 9

Supplemental Datafile 10

Supplemental Datafile 11

## Acknowledgments

We thank Dr. Joanna Rorback for providing the *IF3*-KO cell line. We thank Dr. Mathew F. Allan for his assistance with using DREEM. We thank Ahram Ahn for his assistance with AUROC calculations.

This research was supported by National Institute of General Medicine (NIGMS) grant R35-GM118141 (to A.Ba.), Chan Zuckerberg Initiative Neurodegeneration Challenge Network Collaborative Pairs Pilot Project Award (to A.Ba. and S.R.), the Burroughs Wellcome Fund ID 1018246.01 (S.R. and A.Ba.), and the US Department of Defense grant PR180598 (to F.F.). J.C.M was supported by the Eunice Kennedy Shriver National Institute of Child Health & Human Development of the National Institutes of Health under Award Number F30HD107939.

## Author contributions

A.Ba., S.R., F.F., and J.C.M. conceived the study; J.C.M., F.F., and M.B. performed in organello assays and analysis of mitochondrial translation; A.Bri. and S.R. performed in vitro assays and analysis of most DMS-MaPSeq using DREEM; A.Bri., J.C.M., M.B., and A.B. prepared panels and figures; all authors contributed to data interpretation; A.B. wrote the first draft of the paper. All authors edited the manuscript. All authors read and approved the final manuscript.

## STAR Methods

### Resource Availability

#### Lead contact

Further information and requests for resources and reagents should be directed to and will be fulfilled by the Lead Contact, Dr. Antoni Barrientos, Ph.D.

#### Materials availability

All unique/stable reagents generated in this study (plasmids and cell lines) are available from the Lead Contact with a completed Materials Transfer Agreement.

#### Data and code availability

All Source Data is either included in the manuscript or will be provided upon request. Sequencing data have been uploaded to Gene Expression Omnibus (GEO), with accession number GSE246523.

#### Experimental Models

##### Human cell lines and cell culture conditions

Human HEK293T embryonic kidney cells (CRL-3216, RRID: CVCL-0063) were obtained from ATCC. HEK293T *LRPPRC*-KO ^45^ and *IF3*-KO ^42^ cell lines were previously reported. Cells were cultured in high-glucose Dulbecco’s modified Eagle’s medium (DMEM, Life Technologies) supplemented with 10% fetal bovine serum (FBS), 5 mL 100X antibiotic/antimycotic, 1 mM sodium pyruvate, 1 mM sodium formate, and 50 g/ml uridine, and 1x GlutaMAX (ThermoFisher Scientific; Waltham, MA) at 37°C under 5% CO2. Cell lines were routinely analyzed for mycoplasma contamination.

##### Key reagents

Tables presenting the list of antibodies, recombinant DNAs, oligonucleotides, and siRNA oligoribonucleotides used in this study are included in **Key Resources Table**.

### Methods Details

#### The mitochondrial DMS-MaPseq approach

##### Mitochondrial Isolation and in organello DMS labeling

One or five T-175 flasks of wild-type or *LRPPRC-*KO cells, respectively, grown at 80% confluency, were inoculated into one liter of Freestyle 293 expression medium supplemented with 50 mL FBS, 1 mM sodium formate, 50 μg/ml uridine, and 10 mL of 100X antibiotic/antimycotic. Cells were grown at 37°C under 5% CO_2_, spinning at 63 RPM for five days. Mitochondria were isolated as previously described ^78^ and resuspended in 1 mL of STE buffer (0.32 M sucrose, 1 mM EDTA, and 10 mM Tris-HCl, pH 7.4). 3 mg of freshly prepared mitochondria were pelleted by centrifugation at 10,000 *xg* for 10 min at 4 °C. The pellet was resuspended in 150 μL of Na-cacodylate buffer (300 mM Na-cacodylate, 6 mM MgCl_2_, and 0.32 M sucrose) that had been preheated to 37 °C and incubated for 3 min at 37 °C. A freshly prepared and preheated mixture of 150 μL of the Na-cacodylate buffer plus 3 μL of DMS was added to the mitochondria, resulting in a final DMS concentration of 1%. Mitochondria were incubated for 5 min at 37 °C with gentle shaking. To quench the DMS reaction, 180 μL of beta-mercaptoethanol was added to the mitochondria, the solution was mixed by pipetting up and down several times, and the sample was centrifuged at 10,000 *xg* for 10 min at 4 °C. The mitochondrial pellet was washed twice in 1 mL of a glycerol buffer (10% glycerol, 0.15 mM MgCl_2_, 10 mM Tris-HCl pH 6.8), following the same centrifugation conditions indicated above and resuspended in 1 mL of Trizol.

##### Total RNA extraction and DNase treatment

RNA was extracted following the Trizol manufacturer’s specifications. The aqueous phase was transferred to a new tube, and an equal volume of 100% isopropanol and 3 μL of glycogen were added to precipitate the RNA. The samples were incubated at −80 °C overnight and centrifuged at 15,000 *xg* for 45 min, at 4 °C. RNA was resuspended in 50 μL of RNAse-free water and quantified by measuring absorbance at a wavelength of 260 nm with Nanodrop (ThermoFisher Scientific). 10 μg of RNA was DNase-treated using the TURBO DNase kit (ThermoFisher Scientific). After treatment, the RNA was purified using RNA Clean and Concentrator −5 kit (Zymo) according to the manufacturer’s protocol. DNase treatment and column purification were repeated two additional times to ensure the complete removal of all contaminating DNA.

##### rRNA subtraction

We established a custom rRNA subtraction protocol, based on Phelps, et al. 2021 ^32^, to subtract rRNAs, and more specifically, the mitochondrial rRNAs, which represent the more abundant RNA species in the sample. Oligomers against the mitochondrial rRNAs were designed using the Oligo-ASST tool ^32^, and combined with oligomers against cytosolic RNAs ^79^. Specifically, a 10X mix of oligomers was created at a 2:1 mitochondrial:cytosolic rRNA oligomer ratio, resulting in a concentration of 500 nM per mitochondrial oligo, and 250 nM per cytosolic oligo.

One microgram of DNase-treated RNA per reaction was used as the input for the rRNA subtraction (multiple subtraction reactions are required for sufficient input for library preparation). First, 1 μl of the above rRNA subtraction oligo mix and 2 μl 5X hybridization buffer (1X concentration: 200 mM NaCl, 100 mM Tris-HCl, pH 7.4) were added to each reaction, and the final volume was adjusted with RNAse-free water to 10 μl. The samples were denatured at 95 °C for 2 min, and then the temperature was reduced by 0.1 °C/s until the reaction reached 22°C. Next, a mix of 2 μl RNase H buffer, 2 μl thermostable RNase H, and 6 μL of water was added. The samples were incubated at 65°C for 5 min. The DNA oligos were degraded by adding a mix of 2.5 μL TURBO DNase, 5 μL TURBO DNase buffer, and 22.5 μL of water, and incubating at 37°C for 30 min. The RNA was cleaned with RNA Clean and Concentrator −5, following the manufacturer’s instructions for selecting RNA molecules >200 nt, and eluted in 12 μl of RNAse-free water.

##### Library generation

Sequencing libraries were generated using the xGen Broad-range RNA library prep kit from IDT, following the manufacturer’s specifications, with the following modifications. The RNA fragmentation step was conducted by combining 1 μg of RNA with 1 μL of reagent F1, 4 μL of F3, and 2 μL of F4, and incubating at 95 °C for 2 min. The samples were immediately placed on ice and combined with a mixture of 1 μL 10 mM TGIRTIII reverse transcriptase (RT), 1 μL R1 reagent, and 1 μL 0.1 M DTT. This RNA:RT mixture was incubated for 30 min at room temperature before adding 2 μL of reagent F2, and then placed in a thermocycler for 10 min at 20°C, 10 min at 42 °C, and 60 min at 55 °C. The thermocycler was programmed to pause at 55°C, at which point 1 μL of 4M NaOH was added and programmed to continue at 95°C for 3 minutes. At this point, the sample was removed from the thermocycler and placed on ice, and 1 μL of 4M HCl and 27 μL of low EDTA TE were added. After the indexing PCR, the indexed libraries were resolved on an 8% TBE gel for 55 min at 180 V and size-selected for products ~300 nt. The products were sequenced with NovaSeq6000 system (paired-end run,150 cycles).

#### Pulse labeling of mitochondrial translation products

To determine mitochondrial protein synthesis, 6-well plates were pre-coated at 5 μg/cm^2^ with 50 μg/mL collagen in 20 mM acetic acid and seeded with WT or LRPPRC cell lines ^45^. 70 % confluent cell cultures were incubated for 30 min in DMEM without methionine and then supplemented with 100 μl/ml emetine for 10 min to inhibit cytoplasmic protein synthesis as described ^80^. 100 μCi of ^[35^S]-methionine was added and allowed to incorporate into newly synthesized mitochondrial proteins for 15 minutes. Subsequently, whole-cell extracts were prepared by solubilization in RIPA buffer, and equal amounts of total cellular protein were loaded in each lane and separated by SDS-PAGE on a 17.5 % polyacrylamide gel. Gels were transferred to a nitrocellulose membrane and exposed to a Kodak X-OMAT X-ray film. The membranes were then probed with a primary antibody against β-ACTIN as a loading control. Optical densities of the immunoreactive bands were measured using the Histogram function of the Adobe Photoshop software in digitized images.

#### *In organello* translation and immunoprecipitation

Two mg of freshly-isolated mitochondria were pelleted by centrifugation at 8,000xg for 5 minutes at 4°C. Mitochondria were washed with 1 mL of *in organello* translation buffer (ITB: 100 mM Mannitol, 10 mM Na-Succinate, 80 mM KCl, 5 mM MgCl_2_ 1 mM Na-Phosphate, 25 mM HEPES, pH 7.4, 60 μg/mL of each of the 20 amino acids (no methionine), 5 mM ATP, 20 μM GTP, 6 mM creatine phosphate, and 60 μg/mL creatine kinase) and spun again at the same speed. Mitochondria were resuspended in 1 mL of ITB preheated to 37 °C and incubated for five minutes at 37°C. Mitochondria were then pulsed with 100 μCi of [^35^S]-methionine for 15 and 30 minutes at 37°C and centrifuged at 8,000 *xg* for 5 minutes at 4°C. Mitochondria were resuspended in 200 μL of lysis buffer (150 mM KCl, 20 mM HEPES, pH 7.4, 5 mM EDTA, 0.5 mM PMSF, and 1% Lauryl Maltoside (LM)), incubated on ice for five minutes, and centrifuged at 22,000xg for 15 minutes at 4°C to clarify the lysate. The supernatant was diluted by the addition of 200 μL of lysis buffer without LM and incubated for 4 h at 4 °C with constant gentle shaking with 50 μL of either mouse IgG-agarose beads (Sigma A0919), protein A agarose beads (ThermoFisher 20334) plus 8 uL of MT-ATP8 N-terminal antibody (Invitrogen, PA5-75605), or protein A agarose beads plus 8 uL of MT-ATP8 C-terminal antibody (Cell Signaling, 174125). After incubation, the beads were pelleted by spinning at 200xg for 2 minutes, and the supernatant was removed and saved as the unbound fraction. The beads were then washed 3x with 1 mL PBS, spinning at 200 *xg* for 2 min after each wash. Beads were then incubated with 100 μL of 1x Laemmli Buffer^81^ for 15 min with gentle shaking at room temperature to elute the bound proteins. 20 μL of the unbound samples and 100 μL of the bound samples were loaded onto a 12-20 % gradient polyacrylamide gel, run at 110 V overnight, transferred to a nitrocellulose membrane, and exposed to a Kodak X-OMAT X-ray film.

For chase experiments, samples were pulsed for 30 minutes with 100 μCi of [^35^S]-methionine and spun down as described above. Samples were then resuspended in 1 mL of ITB supplemented with 60 μg/mL of non-radioactive methionine and incubated for 1 or 2 h at 37 °C. Following the chase period, samples were lysed and processed for immunoprecipitation as described above.

#### *In vitro* transcribed mitochondrial mRNA and DMS modification

gBlocks were obtained from IDT for the mature mt-mRNAs and the 7S RNA (see Key Resources for sequences). T7 promoters were added by PCR, and used for T7 Megascript in vitro transcription (ThermoFisher Scientific) according to the manufacturer’s instructions with a 16 h incubation time at 37 °C. Subsequently, 1 μl Turbo DNase I (ThermoFisher Scientific) was added to the reaction and incubated at 37 °C for 15 min. The RNA was purified using RNA Clean and Concentrator −5 kit (Zymo).

For DMS modification, 10 μg of RNA in 10 μl H_2_O was denatured at 95 °C for 1 min and then placed on ice. Based on the DMS concentration used in the next step, 300 mM sodium cacodylate buffer (Electron Microscopy Sciences) with 6 mM MgCl_2_ (refolding buffer) was added so that the final volume was 100 μl. (*e.g*., for 2.5 % final DMS concentration: add 87.5 μl refolding buffer and 2.5 μl DMS) Then, 2.5 μl was added and incubated at 37 °C for 5 min while shaking at 500 r.p.m. on a thermomixer. The DMS was neutralized by adding 60 μl β-mercaptoethanol (Millipore-Sigma). The RNA was purified using RNA Clean and Concentrator −5 kit.

The library generation protocol, as described above, was used after this step.

##### Mapping and Bit Vector Generation

Trimmed FASTQ files were aligned to the human mitochondrial genome (NCBI: NC_012920.1) using Bowtie2 ^82^ with the following parameters: --local --no-unal --no-discordant --no-mixed -L 12 -X 1000. Only reads aligned exactly one time were considered for further analysis. Mapped reads were then classified as originating from the heavy (sense) strand or light (antisense) strand of the mitochondrial genome using a custom Python script ^37^. Read pairs were classified as antisense if they had the following SAM flag ^83^: PAIRED and PROPER_PAIR and ({READ1 and MREVERSE and not REVERSE} or {READ2 and REVERSE and not MREVERSE}) and not (UNMAP or MUNMAP or SECONDARY or QCFAIL or DUP or SUPPLEMENTARY).

Bit vectors were generated from the resulting stranded SAM files as described in Tomezsko et al. 2019 ^31^. Briefly, a bit vector the size of the reference genome was generated for each set of paired-end reads in the stranded SAM files. For each position where the read matched the reference sequence, the bit vector value was set to 0, and for each position at which there was a mismatch or deletion, the value was set to 1. Any position with a Phred score < 20 or no coverage over the reference genome was labeled “uninformative” and not considered. For each position along the reference genome, the total coverage over that position, the number of 0s, and the number of 1s, as calculated using DREEM ^31^. Population average DMS reactivities were then calculated by taking the ratio of the number of mismatches and deletions to total coverage at individual positions. Any positions with DMS reactivity > 0.3 were excluded from downstream analysis and removed from the population average.

##### Genome-wide bivector coverage, signal-to-noise calculation

Genome-wide bit vector coverage (**Fig. 1B**) was determined by counting the number of 0s or 1s in all bit vectors at each position in the mitochondrial genome. The genome-wide signal (**Fig. 1C**) was calculated by taking the mean DMS reactivity of As and Cs every 80nt, starting at nucleotide 41 and advancing by a single nucleotide at each window step. Genome-wide “noise” was calculated in the same manner, except the mean DMS reactivity was calculated on Gs and Us.

##### Genome-wide Gini Index and Correlation calculations

Sliding window genome-wide Gini index was calculated over 80nt windows of the DMS reactivities As and Cs from the population average, starting at position 41 and advancing by a single nucleotide for each window step.

Genome-wide Pearson correlation of DMS reactivities was calculated between biological replicate population average reactivities (**Fig. 3A**) and between the reactivity profiles of the wild-type (WT) and *LRPPRC* knockout (KO) conditions, as depicted in Fig. 3a and Fig. 3b. This correlation analysis was over 80nt sliding windows of As and Cs. Positions for which both datasets did not have reactivity data were excluded from the calculation. The position of the correlation values represents the center of the sliding window.

#### *rRNA* agreement and area under the curve calculations

The secondary structure of the human mitochondrial 12S ribosomal RNA was downloaded from Amunts *et al*, 2015 ^35^. Paired bases in the secondary structure model were labeled “True” while unpaired bases were labeled “False.” To determine how well our population average DMS reactivities agreed with the “ground truth” rRNA structure, ROC curves and AUROC values were calculated with the R package ‘pROC’ ^84^. Only DMS reactivities on As and Cs were considered. “True Positives” in the ROC curve represent paired nucleotides with DMS reactivities less than the sliding threshold, while “False positives” represent unpaired nucleotides with DMS reactivities less than the sliding threshold.

##### DMS Normalization

When indicated, DMS reactivities were normalized to a scale of 0-1. DMS reactivities on Gs and Us were set to 0. The median DMS reactivity of the top 10% most reactive DMS positions was calculated and used to divide all DMS reactivities. Any remaining positions with a normalized reactivity > 1 were winsorized ^85^ by setting their normalized DMS reactivity to 1.

##### Folding mitochondrial mRNAs

Population average DMS reactivities for individual mitochondrial mRNAs were normalized as previously described. Any position without DMS reactivity values, including Gs and Us, was set to −999. Entire mRNAs were then folded using the “Fold” command from RNAstructure ^34^ with the following parameters: -m 3 to generate three structures, -md 350 to specify a maximum pairing distance of 350 nt, -dms to use the normalized DMS reactivities as constraints on the folding algorithm.

##### Ribosome Profiling Normalization

Alignment files for MitoRibosome profiling were sourced from Pearce et al. 2017 ^9^. MitoRibosome occupancy on mRNAs was determined by extracting read depth from the alignment file using the “depth” command in SAMtools ^83^, with the “-a” option to ensure output at all positions. The depth file was subset to include positions intersecting with mitochondrial mRNAs of interest. In the context of analyzing MitoRibosome occupancy at the *COX1* locus, the initial 40 nucleotides were omitted to exclude a peak associated with mitoribosome initiation. Normalization of subsequent depth values to a 0-1 scale was achieved by determining the maximum read depth value and dividing all other read depth values by that maximum.

##### LRPPRC CLIP Normalization

We utilized existing PAR-CLIP data on LRPPRC, as generated and processed by Siira et al. 2017 ^23^. Given the occurrence of high signal at particular CLIP peaks, especially on mitochondrial rRNAs, and the desire to visualize LRPPRC binding sites on mitochondrial mRNAs, we opted to set an upper limit for displayed LRPPRC depth at 5,000 reads. Consequently, we presented the density of reads emerging from the PAR-CLIP on a scale spanning from 0 to 5,000.

##### Genome-wide clustering of mitochondrial mRNAs

To facilitate clustering analysis, FASTQ files were mapped to individual transcript reference files in the same fashion as previously described. Bit vectors were then generated over 100nt, non-overlapping windows of each mRNA, except for the final window, which varied in length between transcripts but was always shorter than 100nt. Before clustering, the resulting population average bit vectors were then filtered as previously described ^31^. Briefly, bit vectors were discarded if they contained two mutations within 4-nucleotides, contained a mutation adjacent to an uninformative bit, or possessed too many mutations such that the number of mutated positions exceeds both 10% of the bit vector length and three standard deviations above the median number of mutations across all bit vectors. Additionally, bit-vectors were discarded if they overlapped with less than 20% of the windowed reference sequence. These filtered bit vectors were then clustered using DREEM ^31^ with K= 3; however, we limited our analysis to K = 2 clusters.

The resulting clusters were deemed invalid and discarded for subsequent analysis if the maximum DMS reactivity was greater than 0.3, if the ratio of the maximum DMS reactivity in cluster 1 and cluster 2 was less than 1/3 or greater than 3, or if the number of bitvectors passing filters was less than 100,000. Gini indices and Pearson correlation were then calculated on As and Cs from each valid cluster window individually.

#### Visualizing RNA structures

RNA secondary structure plots were generated in VARNA ^86^ with the color corresponding to normalized DMS values. Arc plots were generated using the R4RNA ^87^ package in R.

#### Accuracy of secondary structures based on average DMS reactivity data and area under the curve calculations

To assess the accuracy of the predicted structures based on average population DMS reactivity data, we employed AUROC, as reported ^37^. AUROC was calculated over sliding windows of 100 nt in 1 nt increments; *x* values represent the centers of the windows. Paired bases in the secondary structure model were labeled “True” while unpaired bases were labeled “False.” To determine how well our population average DMS reactivities agreed with the modeled mRNA structures, ROC curves and AUROC values were calculated with the R package ‘pROC’ ^84^. Only DMS reactivities on As and Cs were considered. “True Positives” in the ROC curve represent paired nucleotides with DMS reactivities less than the sliding threshold, while “False positives” represent unpaired nucleotides with DMS reactivities less than the sliding threshold.

## Supplemental Information

The supplemental information includes 7 supplemental figures and the Key Resources Table.

