## Supplemental material for "The human mitochondrial mRNA structurome reveals mechanisms of gene expression"

### Tables

#### Table 1. Key Resources

| REAGENT or RESOURCE | SOURCE | IDENTIFIER |
|---|---|---|
| Antibodies | | |
| MT-ATP8 N-terminal<br>(use for IP) | Invitrogen | PA5-75605<br>RRID: AB_2719333 |
| MT-ATP8 C-terminal<br>(use for IP) | Cell Signaling | 17412<br>Tested by the manufacturer by immunoblotting using ethidium bromide-treated cells depleted of mitochondrial DNA (mtDNA) and mtDNA-encoded proteins as a negative control |
| Bacterial and virus strains | | |
| NEB 5-alpha competent bacteria | NEB | C2987H |
| Chemicals, peptides, and recombinant proteins | | |
| DMEM, high glucose, pyruvate | Thermo Fisher | 11995073 |
| DMEM, high glucose, no glutamine, no methionine, no cystine | Thermo Fisher | 21013024 |
| Freestyle™ 293 Expression Medium | Thermo Fisher | 12-338-026 |
| DPBS, no calcium, no magnesium | Thermo Fisher | 14190144 |
| Fetal bovine serum | Sigma | 12303C |
| Uridine | Sigma | 58968 |
| Sodium formate | Sigma | 247596 |
| GlutaMAX™ Supplement | Thermo Fisher | 35050061 |
| Collagen I, Rat Tail | Thermo Fisher | A1048301 |
| Complete EDTA Free Protease Inhibitor Cocktail | Sigma | 11873580001 |
| EasyTag™ L-[$^{35}$S]-Methionine | Perkin-Elmer | NEG709A |
| Emetine dihydrochloride | ChemCruz | SC-202600 |
| PMSF protease inhibitor | Thermo | 36978 |
| TIGRTIII | Ingex | TGIRT10 |
| Creatine Phosphate | Sigma | 2380 |
| Creatine phosphokinase | Sigma | C7886 |
| Lauryl Maltoside | Sigma | D4641 |
| Mouse IgG-agarose beads | Sigma | A0919 |
| Protein A agarose beads | Thermo Fisher | 20365 |
| Trizol | Thermo Fisher | 15596026 |
| Critical commercial assays | | |
| TURBO DNase Kit | Thermo Fisher | AM2238 |
| MEGAscript™ T7 Transcription Kit | Thermo Fisher | AM1334 |
| Zymo RNA Clean and Concentrate -5 | Zymo | R1013 |
| SPRIselect Bead-Based Reagent | Beckman Coulter | B23318 |
| Thermostable RNase H | NEB | M0523S |
| xGen UDI Primers Plate 2, 8nt | IDT | 10009816 |
| xGen Broad-range RNA library prep kit | IDT | 10009865 |
| Deposited data | | |
| WT mt transcriptome. DMS mutational data. RNAseq. | GEO | Accession number GSE246523 |

| | | |
|---|---|---|
| *LRPPRC*-KO mt transcriptome. DMS mutational data. RNAseq | GEO | Accession number GSE246523 |
| Experimental models: Cell lines | | |
| HEK293T | ATCC | CRL-3216 |
| HEK293T *LRPPRC*-KO | Soto et al., 2023 [1] | |
| HEK293T *IF3*-KO | Remes et al. 2023 [2] | |
| Oligonucleotides | | |
| See separate table | | |
| Recombinant DNA | | |
| Software and algorithms | | |
| Adobe Illustrator 2023 | | N/A |
| Adobe Photoshop 2023 | | |
| ImageJ V1.53 | | https://imagej.nih.gov/ij/ |
| GraphPad Prism | GraphPad Software v. 10.0.3 (217) | N/A |
| R software 4.3.1 | The R Development Core Team | |
| Varna | Darty et al. 2009 [3] | |
| RNAstructure | Mathew, 2014[4] | |
| DREEM clustering algorithm | Tomezsko et al. 2023 [5] | |
| Other | | |
| Kodak X-OMAT X-ray film | Sigma | F1274-50EA |
| eBlot L1 Protein Transfer System | Genscript | L00686 |
| Novex™ TBE Gels, 8% | Thermo Fisher | EC62152BOX |

#### Table 2. List of oligonucleotides

| Oligo Name | Sequence (5'->3') |
|---|---|
| Oligonucleotides used for mitoribo-depletion | |
| mito_ribodeplete_1 | GGAACGGGGATGCTTGCATGTGTAATCTTACTAAGAGCT |
| mito_ribodeplete_2 | TTTGAGCTGCATTGCTGCGTGCTTGATGCTTGTTCCTTTT |
| mito_ribodeplete_3 | TTATTGCTAAAGGTTAATCACTGCTGTTTCCCGTGGGGGT |
| mito_ribodeplete_4 | TCGTGTGACCGCGGTGGCTGGCACGAAATTGACCAACCCT |
| mito_ribodeplete_5 | ATTGGGGAGGGGGTGATCTAAAACACTCTTTACGCCGGCT |
| mito_ribodeplete_6 | AGCCACTTTCGTAGTCTATTTTGTGTCAACTGGAGTTTTT |
| mito_ribodeplete_7 | TTAGGGCTAAGCATAGTGGGGTATCTAATCCCAGTTTGGG |
| mito_ribodeplete_8 | GGTCCTTTGAGTTTTAAGCTGTGGCTCGTAGTGTTCTGGC |
| mito_ribodeplete_9 | GTGAGGTTGATCGGGGTTTATCGATTACAGAACAGGCTCC |
| mito_ribodeplete_10 | ACTTGCGCTTACTTTGTAGCCTTCATCAGGGTTTGCTGAA |
| mito_ribodeplete_11 | GGGGTAGAAAATGTAGCCCATTTCTTGCCACCTCATGGGC |
| mito_ribodeplete_12 | TCCACCTTCGACCCTTAAGTTTCATAAGGGCTATCGTAGT |
| mito_ribodeplete_13 | GGTGTGTACGCGCTTCAGGGCCCTGTTCAACTAAGCACT |
| mito_ribodeplete_14 | TTAAATGTCCTTTGAAGTATACTTGAGGAGGGTGACGGGC |
| mito_ribodeplete_15 | AGTGCACTTTCCAGTACACTTACCATGTTACGACTTGTCT |

| | |
|---|---|
| mito_ribodeplete_16 | AATGGTTTGGCTAAGGTTGTCTGGTAGTAAGGTGGAGTGG |
| mito_ribodeplete_17 | ATCTTTCCCTTGCGGTACTATATCTATTGCGCCAGGTTTC |
| mito_ribodeplete_18 | GCAGAAGGTATAGGGGTTAGTCCTTGCTATATTATGCTT |
| mito_ribodeplete_19 | CGTCTGGTTTCGGGGGTCTTAGCTTTGGCTCTCCTTGCAA |
| mito_ribodeplete_20 | AATCTTCCCACTATTTTGCTACATAGACGGGTGTGCTCT |
| mito_ribodeplete_21 | AAGATTCTATCTTGGACAACCAGCTATCACCAGGCTCGGT |
| mito_ribodeplete_22 | CAGTTAAATTTACAAGGGGATTTAGAGGGTTCTGTGGGCA |
| mito_ribodeplete_23 | ACTCTCTCTACAAGGTTTTTTCCTAGTGTCCAAAGAGCT |
| mito_ribodeplete_24 | TGGGTGTTGAGCTTGAACGCTTTCTTAATTGGTGGCTGCT |
| mito_ribodeplete_25 | TCTATAGGGTGATAGATTGGTCCAATTGGGTGTGAGGAGT |
| mito_ribodeplete_26 | TCTGACGCAGGCTTATGCGGAGGAGAATGTTTTCATGTT |
| mito_ribodeplete_27 | GGTTGATTGTAGATATTGGGCTGTTAATTGTCAGTTCAGT |
| mito_ribodeplete_28 | CTTTTTTTAACCTTTCCTTATGAGCATGCCTGTGTTGGGT |
| mito_ribodeplete_29 | GGTGATGCTAGAGGTGATGTTTTTGGTAAACAGGCGGGGT |
| mito_ribodeplete_30 | CTACCTTTGCACGGTTAGGGTACCGCGGCCGTTAAACATG |
| mito_ribodeplete_31 | GTAAGAGACAGCTGAACCCTCGTGGAGCCATTCATACAGG |
| mito_ribodeplete_32 | CATAGGGTCTTCTCGTCTTGCTGTGTTATGCCCGCCTCTT |
| mito_ribodeplete_33 | TTAATGCAGGTTTGGTAGTTTAGGACCTGTGGGTTTGTTA |
| mito_ribodeplete_34 | GACTGGTGAAGTCTTAGCATGTACTGCTCGGAGGTTGGGT |
| mito_ribodeplete_35 | GCTGTTATCCCTAGGGTAACTTGTTCCGTTGGTCAAGTTA |
| mito_ribodeplete_36 | TCGGGATGTCCTGATCCAACATCGAGGTCGTAAACCCTA |
| mito_ribodeplete_37 | TCCGGTCTGAACTCAGATCACGTAGGACTTTAATCGTTGA |
| mito_ribodeplete_38 | TAGGCCTTATTTCTCTTGTCCTTTCGTACAGGGAGGAATT |
| mito_ribodeplete_39 | TTCTTGGGTGGGTGTGGGTATAATACTAAGTTGAGATGAT |
| Oligonucleotides used for *in vitro* transcription | |
| COX1_T7_Fwd | TAATACGACTCACTATAGGATGTTCGCCGACCG |
| COX1_Rev | TCTAGATTTTATGTATACGGGTTCTTCGAATGTGTGG |
| COX2_T7_Fwd | TAATACGACTCACTATAGGATGGCACATGCA |
| COX2_Rev | CTATAGGGTAAATACGGGCCCTATTTCAAAGA |
| COX3_T7_Fwd | TAATACGACTCACTATAGGATGGCACATGCAGCGCAAGTAGG |
| COX3_Rev | CTATAGGGTAAATACGGGCCCTATTTCAAAGATTTTTAGGGGA |
| ND1_T7_Fwd | TAATACGACTCACTATAGGATACCCATGGCCAACC |
| ND1_Rev | TAGGTTTGAGGGGGAATGCTGGAGA |
| ND2_T7_Fwd | TAATACGACTCACTATAGG ATTAATCCCCTGGCCCAACCCGTCA |
| ND2_Rev | ATAAGATTATTAGTATAAAAGGGGAGATAGGTAGGAGTAGCGTGGTAAGGG |
| ND3_ T7_Fwd | TAATACGACTCACTATAGGATAAACTTCGCCTTAATTTTAATAATCAACACCCTCCT |
| ND3_Rev | ATTCGGTTCAGTCTAATCCTTTTTGTAGTCAC |
| ND44L_ T7_Fwd | TAATACGACTCACTATAGGATGCCCCTCATTTACATAAATAT |
| ND44L_Rev_ | AAGAGGAAAACCCGGTAATGATGTCGG |
| ND5_T7_Fwd | TAATACGACTCACTATAGGATAACCATGCACACTACTATAACCACCCTAACCCT |
| ND5_Rev | TTCTTCCCACTCATCCTAACCCTACTCCTAATCACATAA |
| ND6_T7_Fwd | TAATACGACTCACTATAGGATGATGTATGCTTTGTTTCTGTTGAGTGTGGGT |

| | |
|---|---|
| ND6_Rev | CCTATTCCCCCGAGCAATCTCAATTACAAT |
| ATP68_T7_Fwd | TAATACGACTCACTATAGGAATGCCCCAACTAAATACTACCG |
| ATP68_Rev | TTATGTGTTGTCGTGCAGGTAGAGGCTTACT |
| CYTB_T7_Fwd | TAATACGACTCACTATAGGATAACCATGCACACTACTATAACCACCCTAACCCT |
| CYTB_Rev | AGGCCCATTTGAGTATTTTGTTTTCAATTAGGGAGATAG |
| 7S_T7_Fwd | TAATACGACTCACTATAGGAGATAAAATTTGAAATCTGGTTAGGCTGGTGTTAGGG |
| 7S_Rev | AATAATAACAATTGAATGTCTGCACAGCCACTTTCC |

### Supplemental Figures

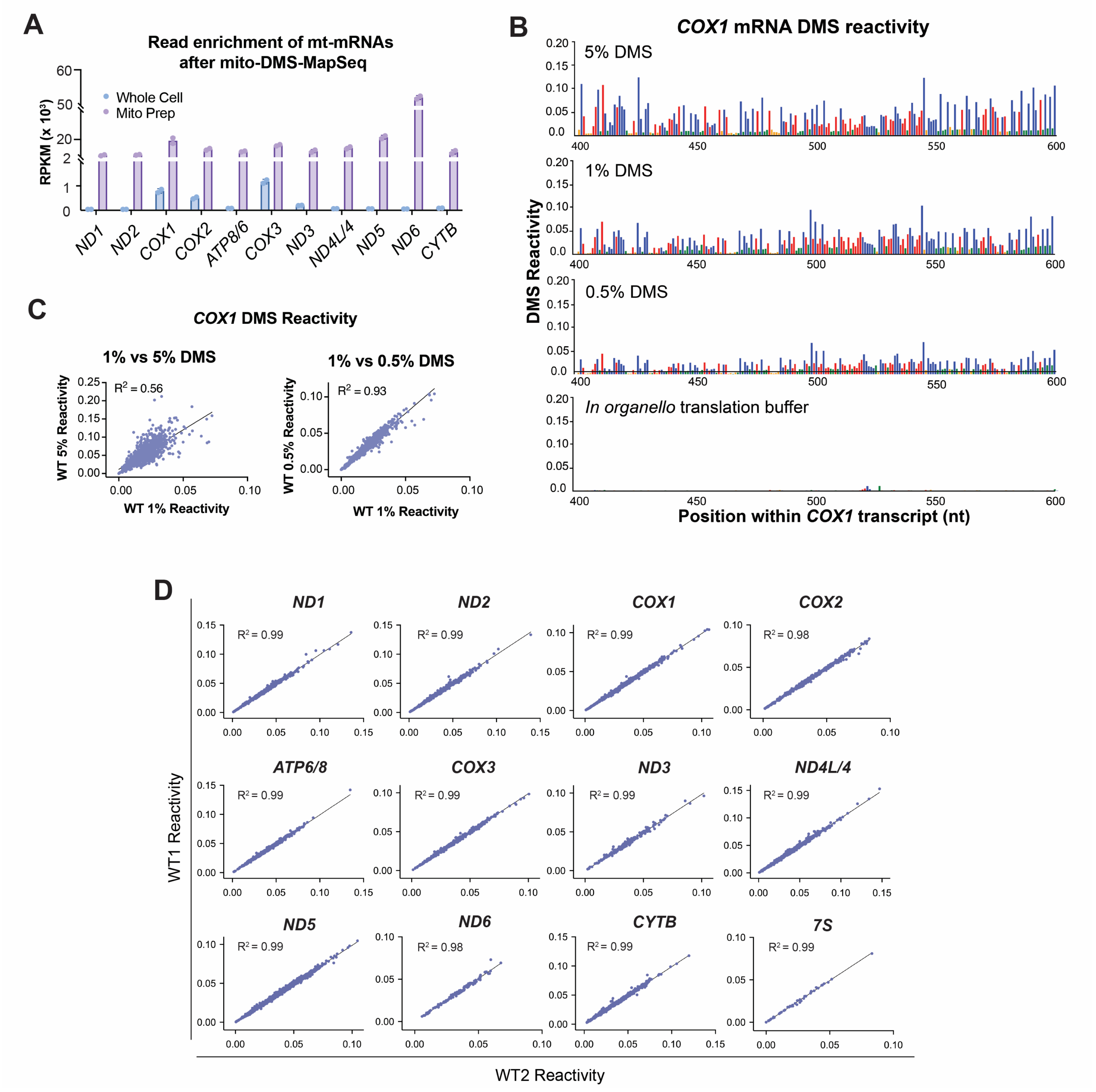

**Supplemental Figure S1. The mito-DMS-MaPseq approach with 1% DMS is reproducible and accurate.** Related to Figure 1.

**(A)** Comparison of enrichment of mitochondrial mRNA reads after DMS-MaPSeq performed in whole cells and in isolated mitochondria.

**(B)** Comparison of DMS reactivities in *COX1* mRNA using 0.1%, 1%, and 5% v/v DMS. A sample with a translation buffer but not DMS was used as a negative control.

**(C)** Correlation of DMS reactivity for *COX1* mRNA between treatment with 1% and 5% DMS, and 1% and 0.5% DMS**.**

**(D)** Interexperimental correlation of DMS reactivity for the indicated 12 mitochondrial transcripts at 1 % v/v DMS. The coefficient of determination $R^2$ values are indicated.

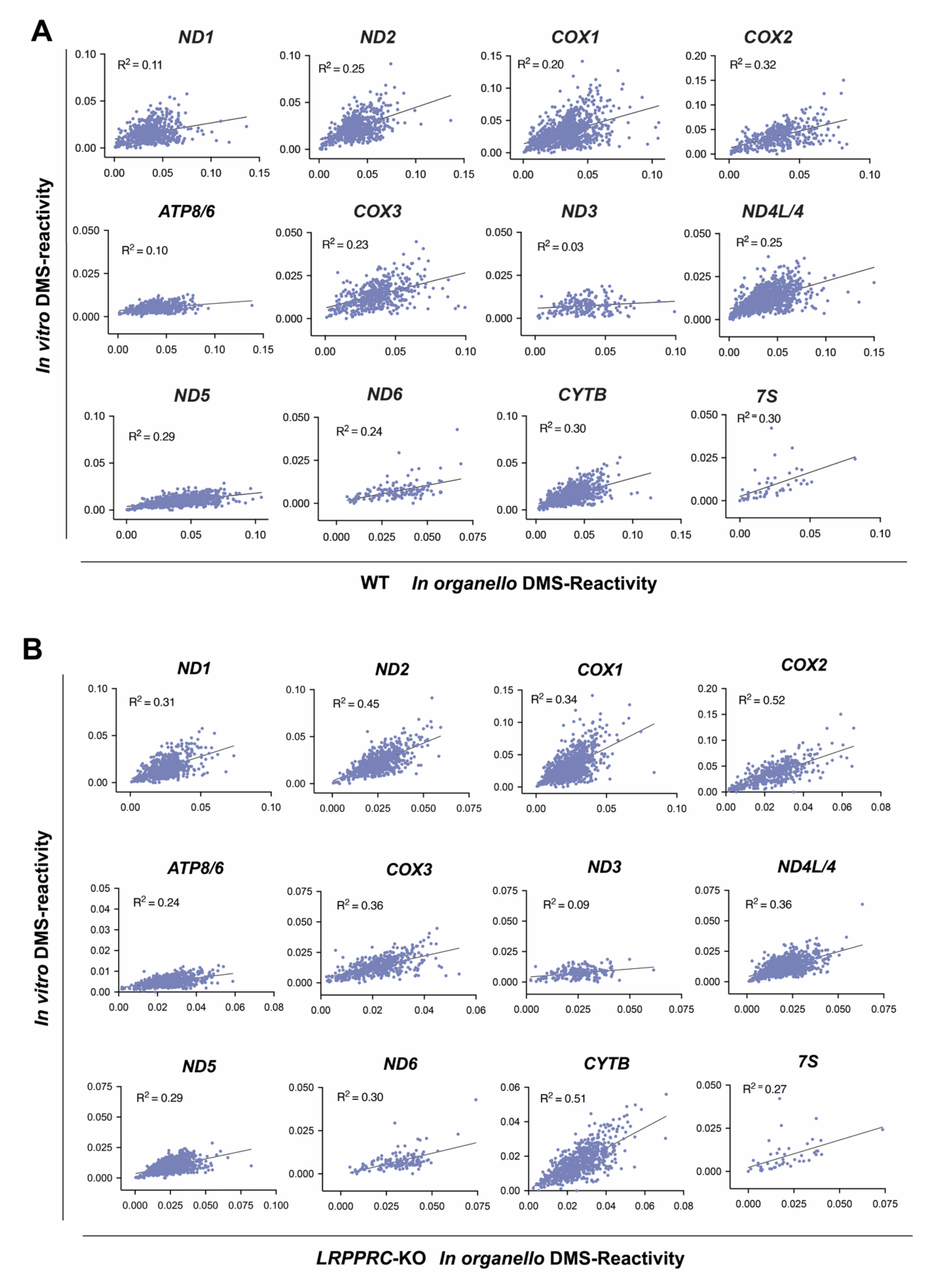

**Supplementary Figure S2. *In organello* structures differ from *in vitro* predictions.** Related to Figure 2.

Correlation of DMS reactivity for the indicated 12 mitochondrial transcripts synthesized and folded natively vs *in vitro* at 1 % v/v DMS. The coefficient of determination $R^2$ values are indicated.

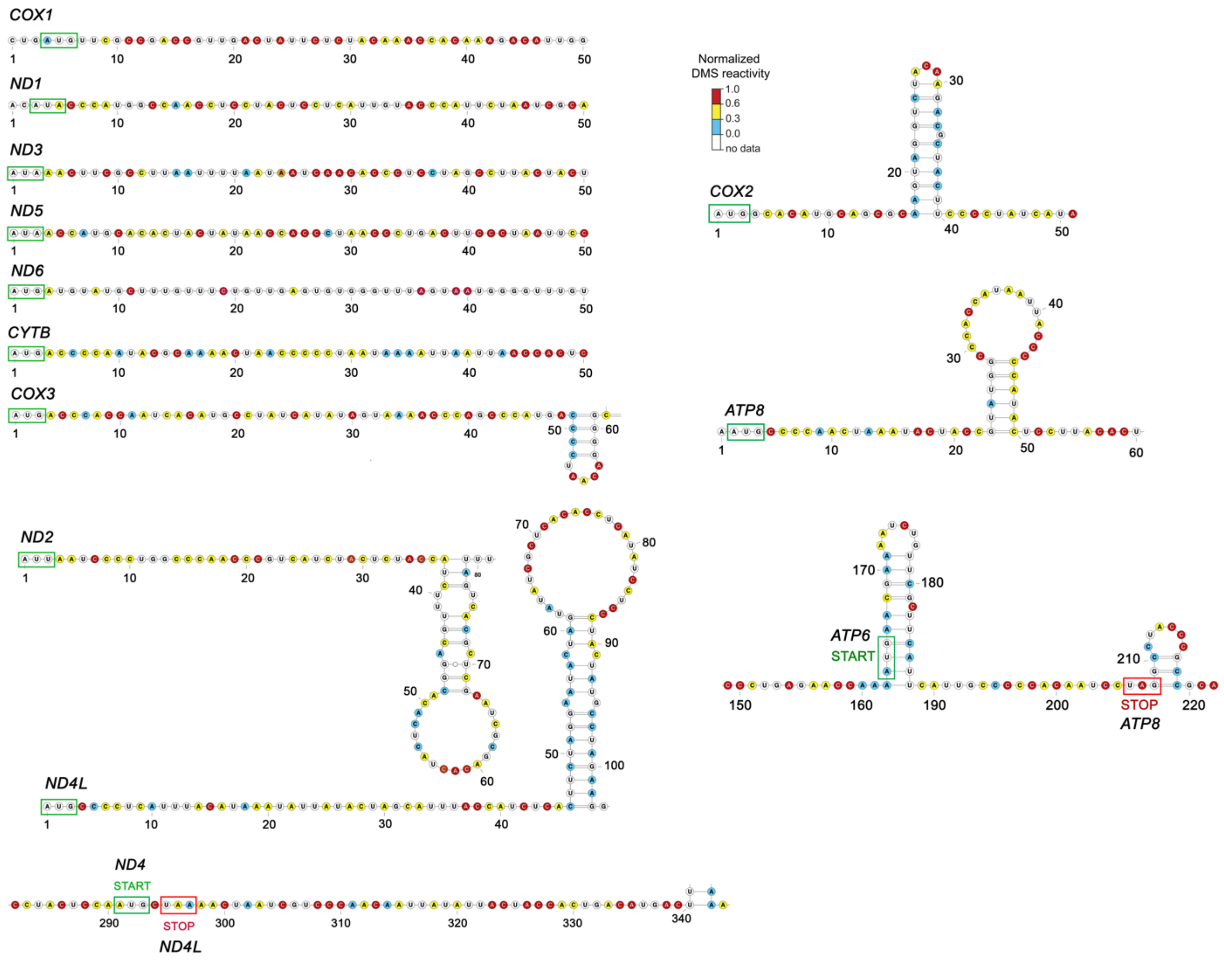

**Supplementary Figure S3. Most 5' regions of mitochondrial mRNAs are unstructured.** Related to Figure 2.

Predicted secondary structure of the ~50 nt downstream of the start codon in the indicated monocistronic and bicistronic transcripts. Nucleotides are colored by normalized DMS reactivity.

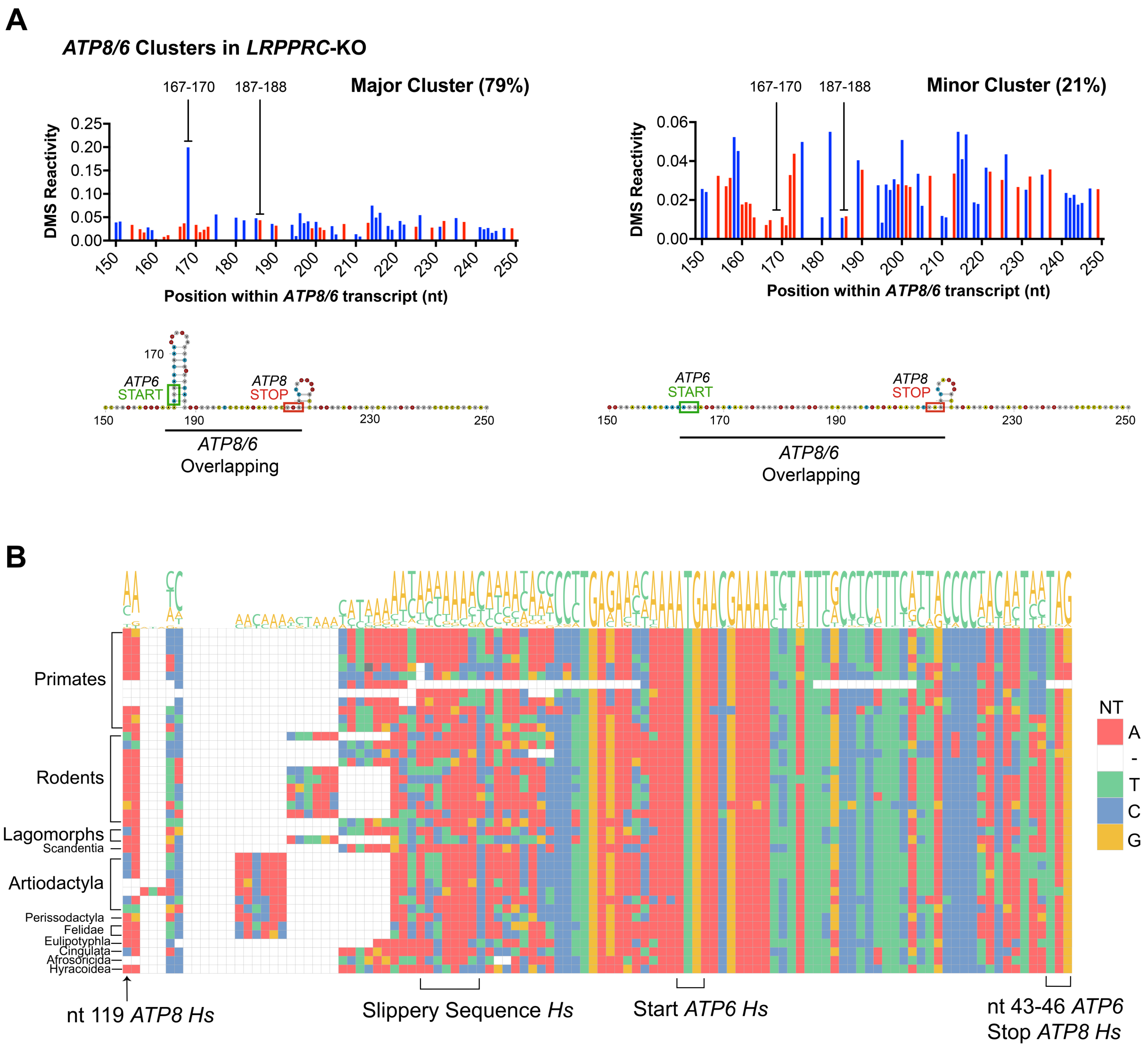

**Supplementary Figure S4. The *ATP8/6* mRNA overlapping region forms a hairpin that is conserved in a major cluster of transcripts in *LRPPRC*-KO mitochondria and whose sequence is conserved across animal species.** Related to Figure 3.

(**A**) DREEM-driven clustering analysis of the mito-DMSMaPseq data on the *ATP8/6* bicistronic transcript in *LRPPRC*-KO mitochondria. For each cluster, the DMS-reactivity plots are shown on the top panel, and the predicted secondary structures are shown on the bottom panel.

(**B**) Alignment of *ATP8/ATP6* sequences from animal mitochondrial genomes. The position of features in the human sequence (*Hs*) is indicated.

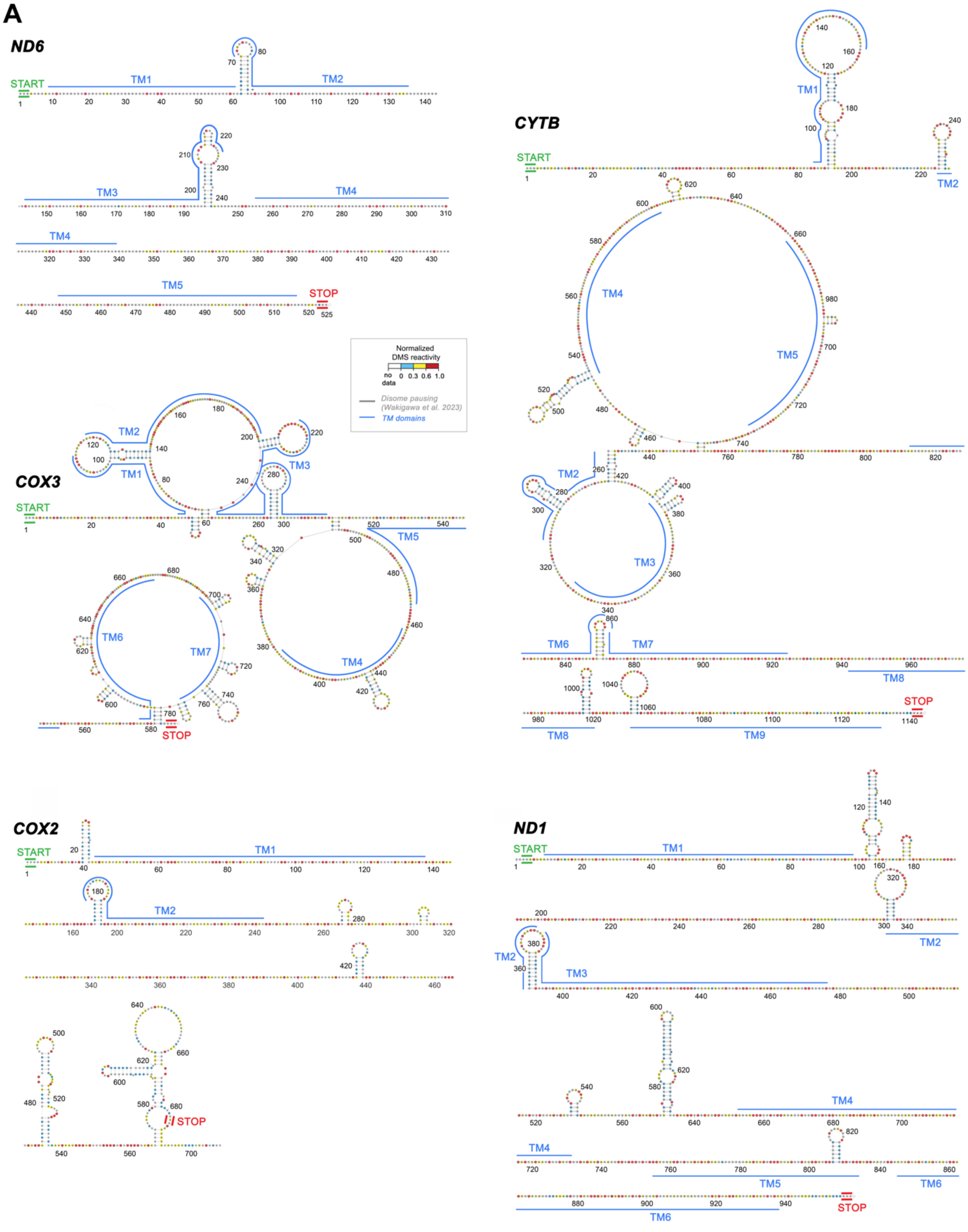
A
ND6
CYTB
COX3
COX2
ND1
START
STOP
TM1
TM2
TM3
TM4
TM5
TM6
TM7
TM8
TM9
Normalized DMS reactivity
no data
0
0.3
0.6
1.0
Disome pausing (Wakigawa et al. 2023)
TM domains

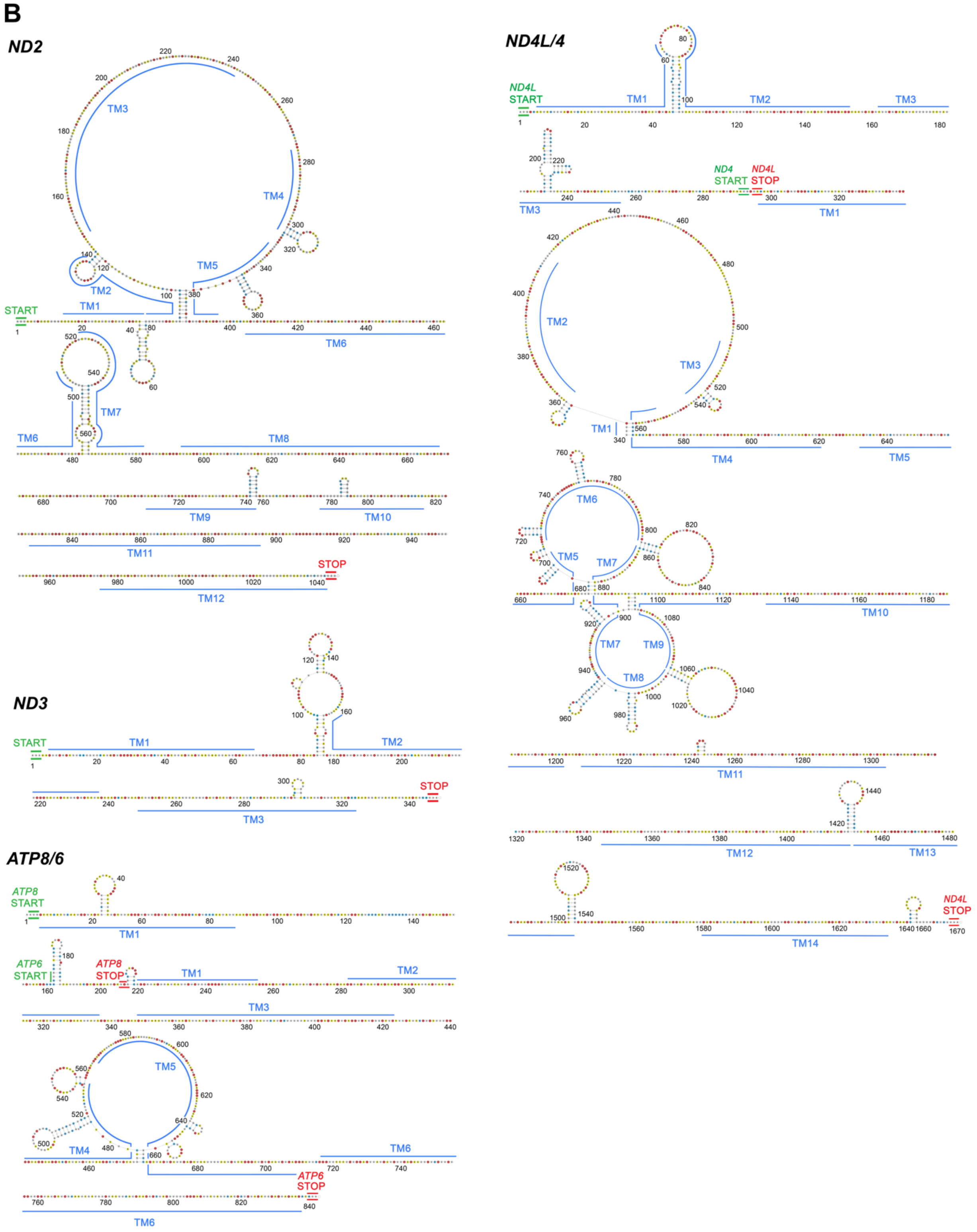

B
ND2
START
TM1
TM2
TM3
TM4
TM5
TM6
TM7
TM8
TM9
TM10
TM11
TM12
STOP
ND3
START
TM1
TM2
TM3
STOP
ATP8/6
ATP8
START
TM1
ATP6
START
ATP8
STOP
TM1
TM2
TM3
TM4
TM5
TM6
ATP6
STOP
TM6
ND4L/4
ND4L
START
TM1
TM2
TM3
TM3
ND4
START
ND4L
STOP
TM1
TM2
TM3
TM1
TM4
TM5
TM6
TM5
TM7
TM10
TM7
TM9
TM8
TM11
TM12
TM13
TM14
ND4L
STOP

C

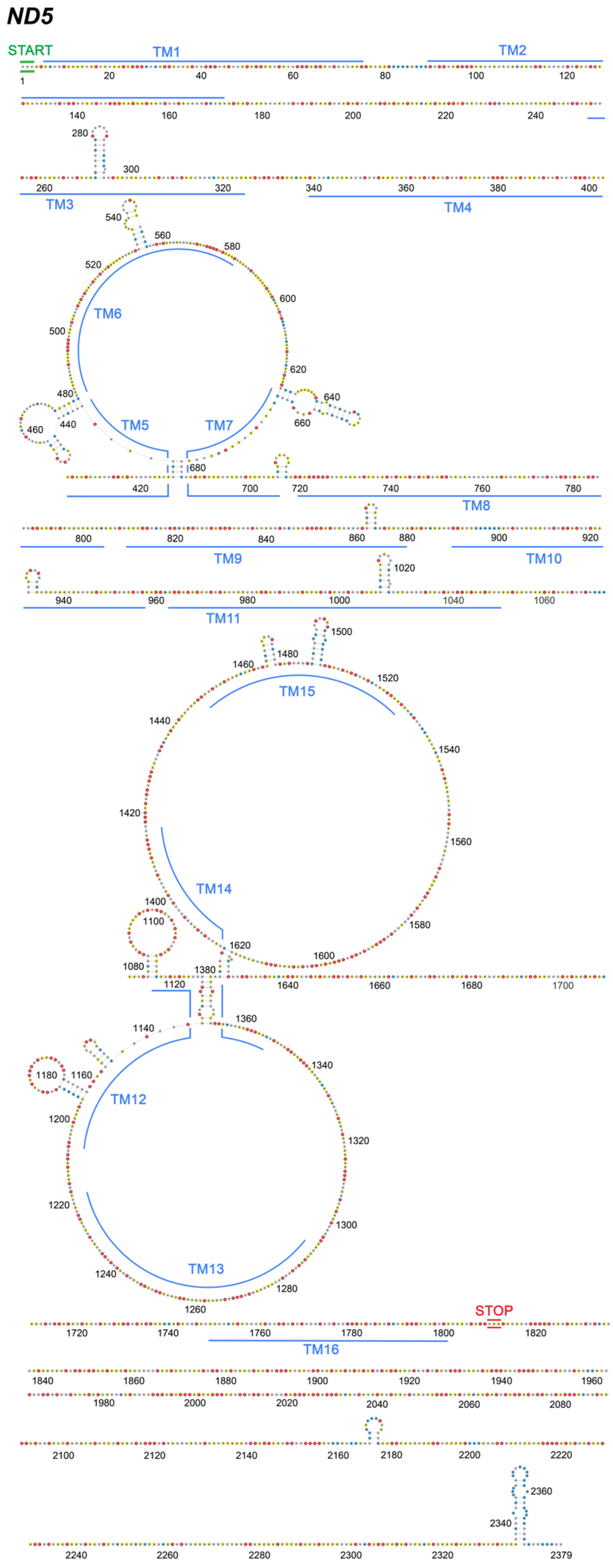

ND5
START
TM1
TM2
TM3
TM4
TM5
TM6
TM7
TM8
TM9
TM10
TM11
TM12
TM13
TM14
TM15
TM16
STOP

**Supplementary Figure S5. Overview of mRNA structure-translation relationships.** Related to Figure 4.

Predicted secondary structure of the indicated transcripts. (**A**) *ND6, COX3, COX2, CYTB* and *ND1* (**B**) *ND2, ND3, ND4L/4*, and *ATP8/6* (**C**) *ND5*. Nucleotides are colored by normalized DMS reactivity. TM domains [6] are indicated with blue lines.

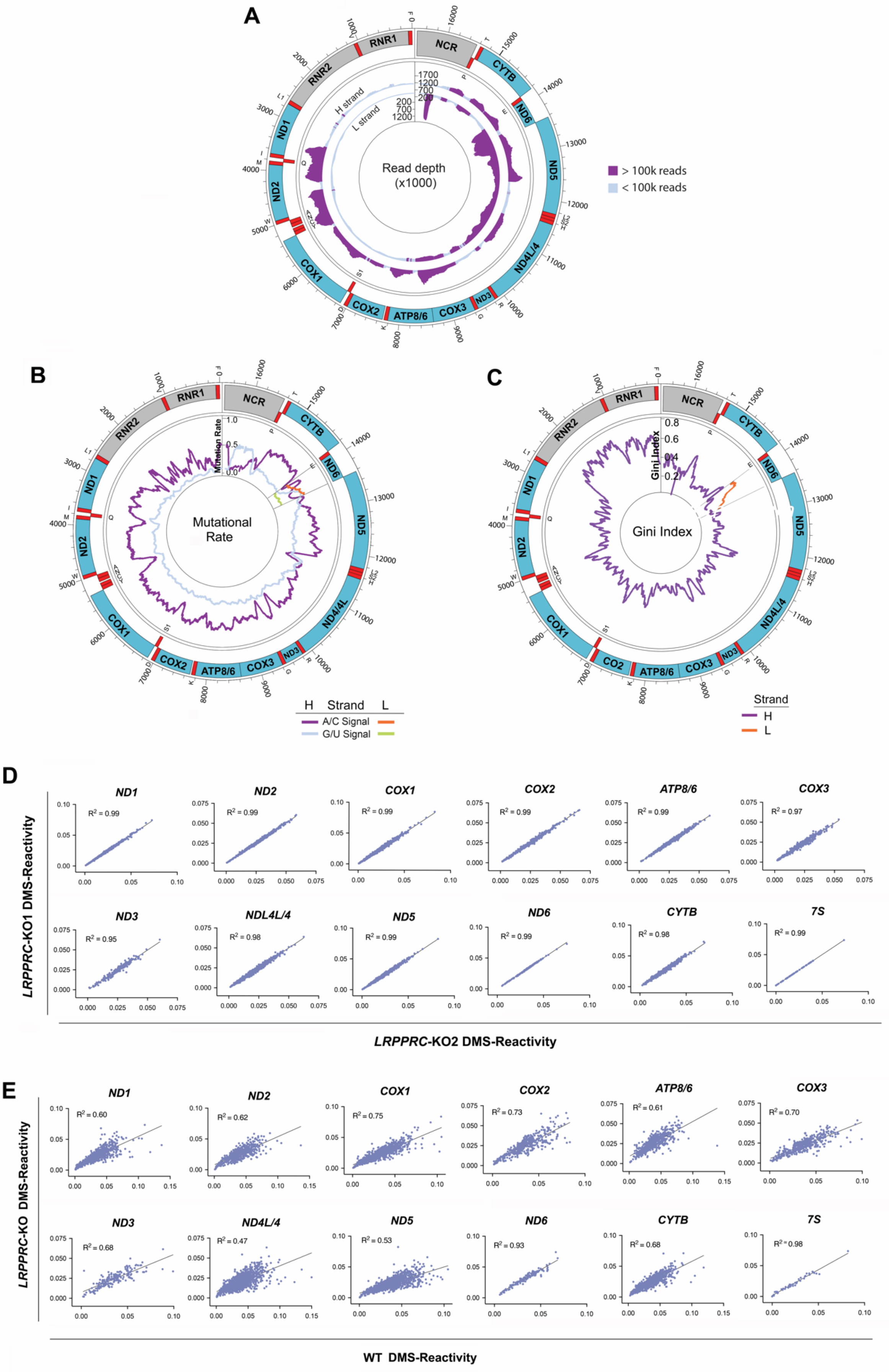
A
Read depth
(x1000)
H strand
L strand
> 100k reads
< 100k reads
RNR1
RNR2
NCR
CYTB
ND6
ND5
ND4L/4
ND3
COX3
ATP8/6
COX2
COX1
ND2
ND1
B
Mutational
Rate
Mutation Rate
H
Strand
L
A/C Signal
G/U Signal
C
Gini Index
Strand
H
L
D
ND1
ND2
COX1
COX2
ATP8/6
COX3
ND3
NDL4L/4
ND5
ND6
CYTB
7S
LRPPRC-KO1 DMS-Reactivity
LRPPRC-KO2 DMS-Reactivity
E
ND1
ND2
COX1
COX2
ATP8/6
COX3
ND3
ND4L/4
ND5
ND6
CYTB
7S
LRPPRC-KO DMS-Reactivity
WT DMS-Reactivity

**Supplementary Figure S6. The Mito-DMS-MapSeq approach reveals a divergent mitochondrial mRNA structurome in *LRPPRC*-KO versus WT mitochondria.** Related to Figure 5.

(**A)** Circular representation of the mitochondrial genome displaying the number of unfiltered bitvectors across the H-strand and L-strand transcriptomes at each position in *LRPPRC*-KO mitochondria.

(**B**) Circular representation of the mitochondrial genome displaying DMS signal/noise ratio (inner tracks) across the heavy (H) and light (L) strands of the mitochondrial transcriptome in *LRPPRC*-KO mitochondria. The purple and light blue lines represent signal *vs*. noise plots of mutation frequencies (*i.e.*, among all reads aligning to each strand coordinate, the fraction of reads with a mutation at that coordinate) on adenines (As) and cytosines (Cs) *vs*. guanines (Gs) and uracils (Us) as a function of strand coordinate for untreated and DMS-treated RNA. A mutation frequency of 0.01 at a given position represents 1% of reads having a mismatch or deletion at that position.

(**C**) Circular representations of the mitochondrial genome H- and L- strands displaying the Gini index of DMS reactivity in *LRPPRC*-KO mitochondria by taking the mean over a sliding window of 100 nt.

(**D**) Interexperimental correlation of DMS reactivity for the indicated mitochondrial transcripts at 1 % v/v DMS revealing high reproducibility. The coefficient of determination $R^2$ value is indicated.

(**E**) Correlation of DMS reactivity for the indicated mitochondrial transcripts in *LRPPRC*-KO vs WT mitochondria. The coefficient of determination $R^2$ value is indicated.

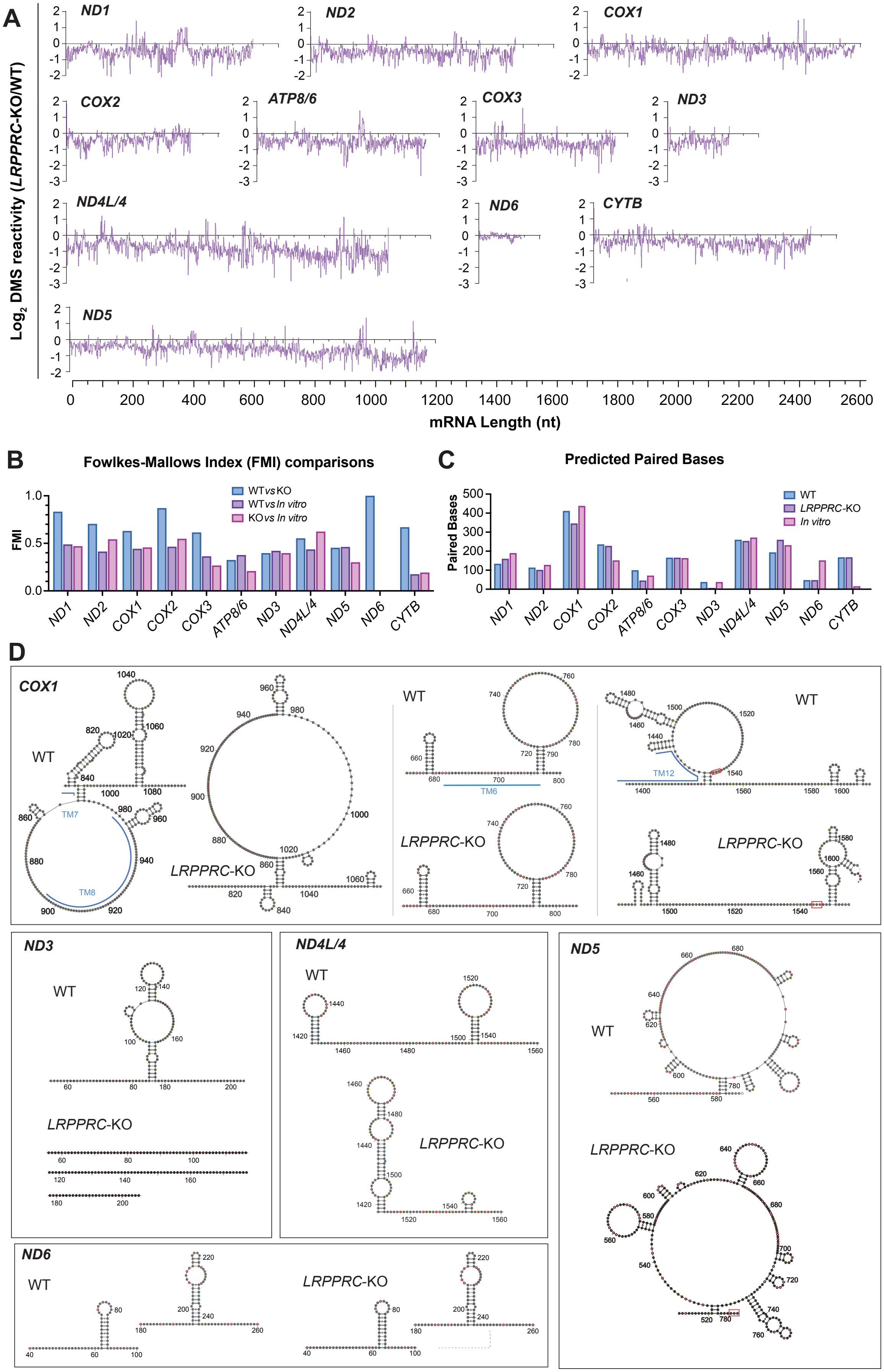

A
Log2 DMS reactivity (LRPPRC-KO/WT)
ND1
ND2
COX1
COX2
ATP8/6
COX3
ND3
ND4L/4
ND6
CYTB
ND5
mRNA Length (nt)
B
Fowlkes-Mallows Index (FMI) comparisons
FMI
WT vs KO
WT vs In vitro
KO vs In vitro
C
Predicted Paired Bases
Paired Bases
WT
LRPPRC-KO
In vitro
D
COX1
WT
LRPPRC-KO
TM7
TM8
TM6
TM12
ND3
ND4L/4
ND5
ND6

**Supplementary Figure S7. The mito-DMS-MapSeq approach reveals a divergent mitochondrial mRNA structurome in *LRPPRC*-KO versus WT mitochondria.** Related to Figure 5.
(**A**) Enrichment of differential mito-DMS-MapSeq counts. Negative values indicate folded regions in *LRPPRC*-KO mitochondria.
(**B**) Predicted transcript structuredness estimated by calculating the number of predicted paired bases normalized by transcript nucleotide length.
(**C**) Comparison of mRNA structures from *LRPPRC*-KO an WT mitochondria using sensitivity and positive predictive value (PPV)[7], whose mean is the Fowlkes-Mallows index (FMI)[8].
(**D**) Comparison of the predicted secondary structure of selected fragments from the indicated transcripts from *LRPPRC*-KO and WT mitochondria. Nucleotides are colored by normalized DMS reactivity. TM domains [6] are indicated with blue lines.
