## Supplementary figures and images for "The human mitochondrial mRNA structurome reveals mechanisms of gene expression"

### Supplemental Datafile 1

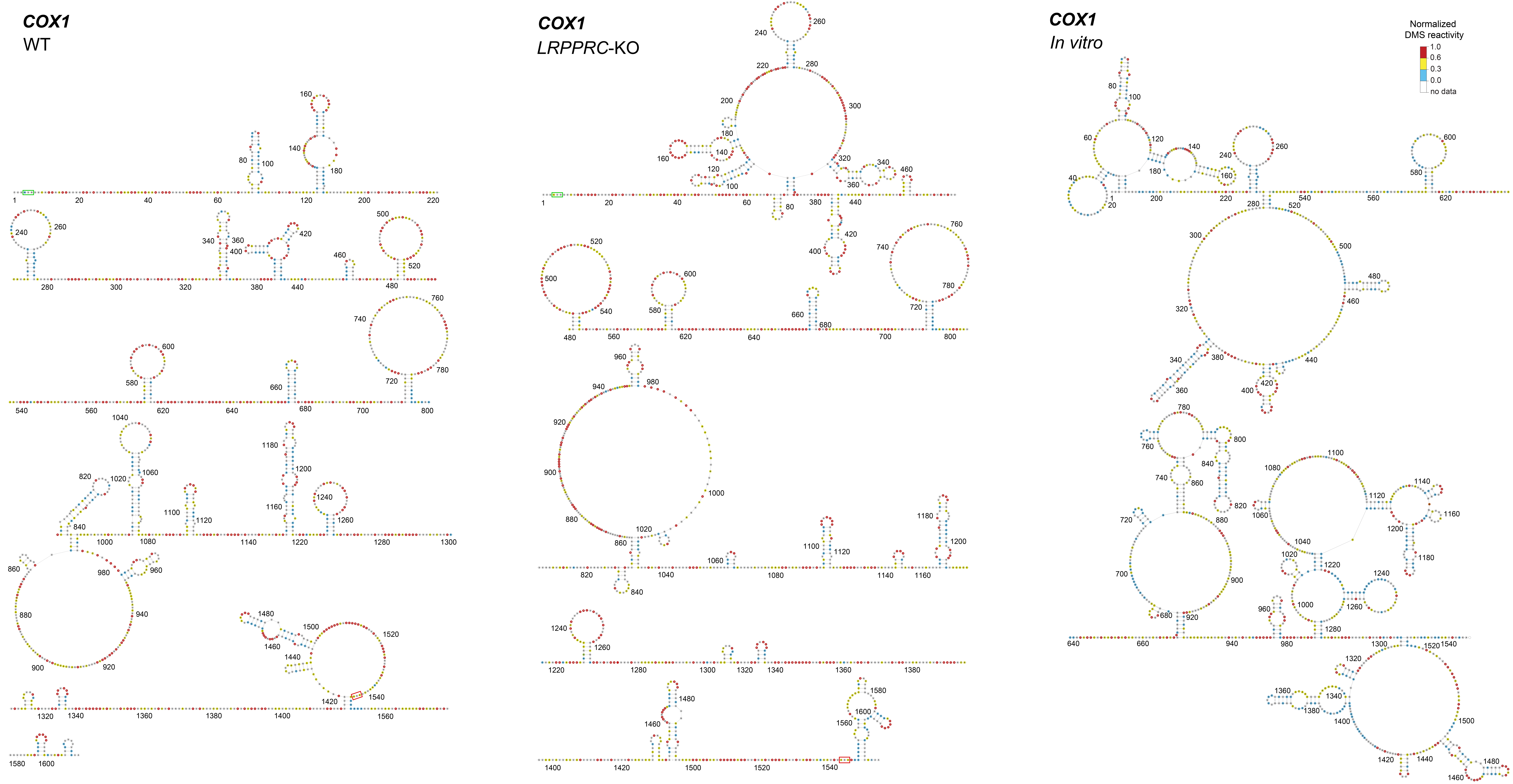

### Supplemental Datafile 2

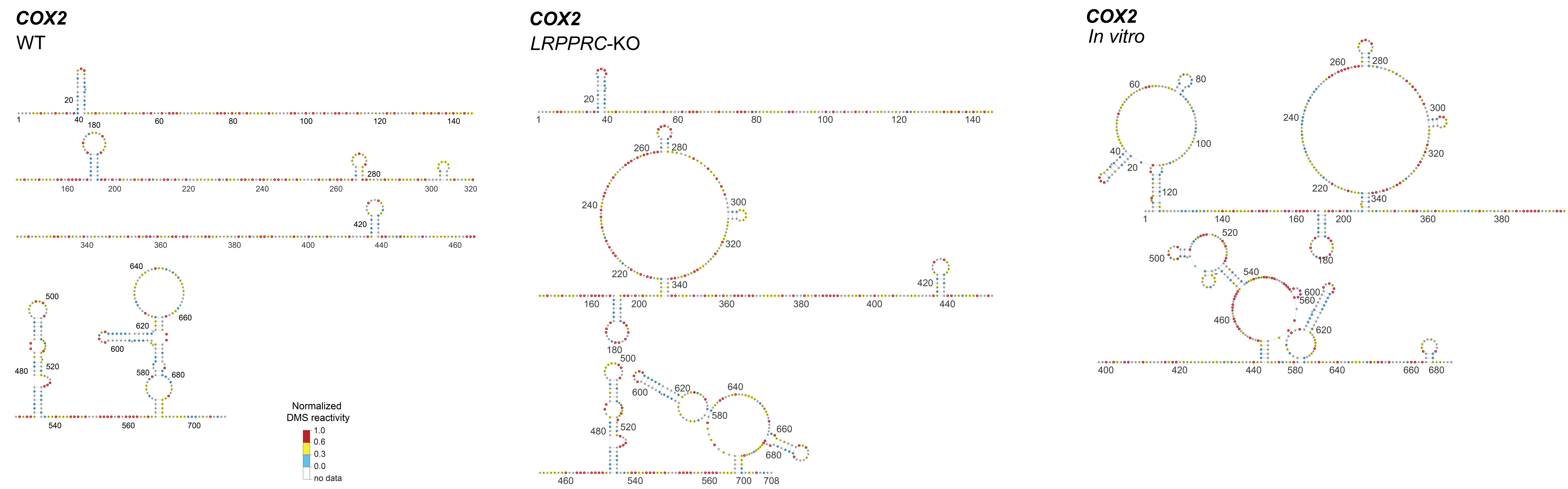

### Supplemental Datafile 3

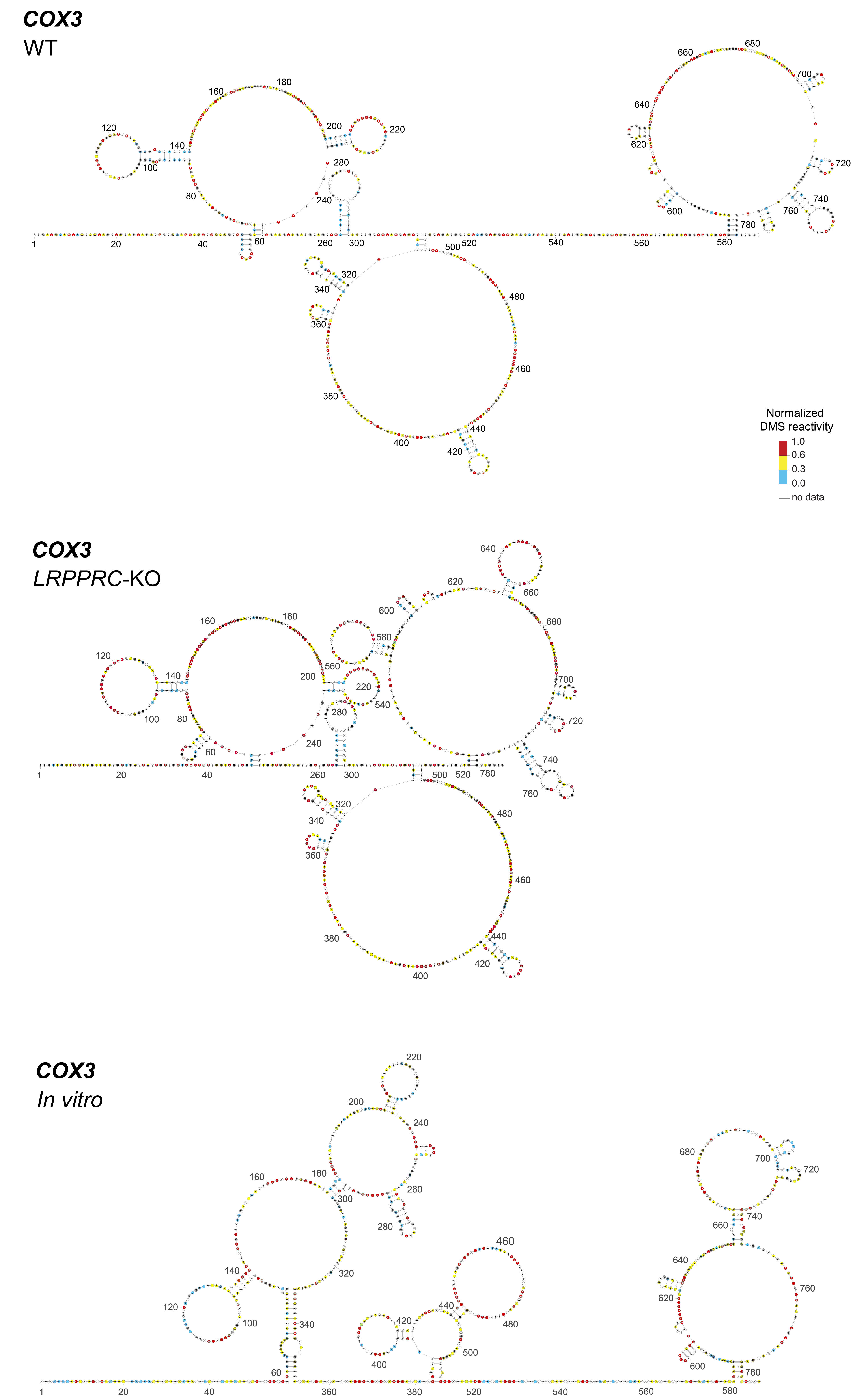

### Supplemental Datafile 4

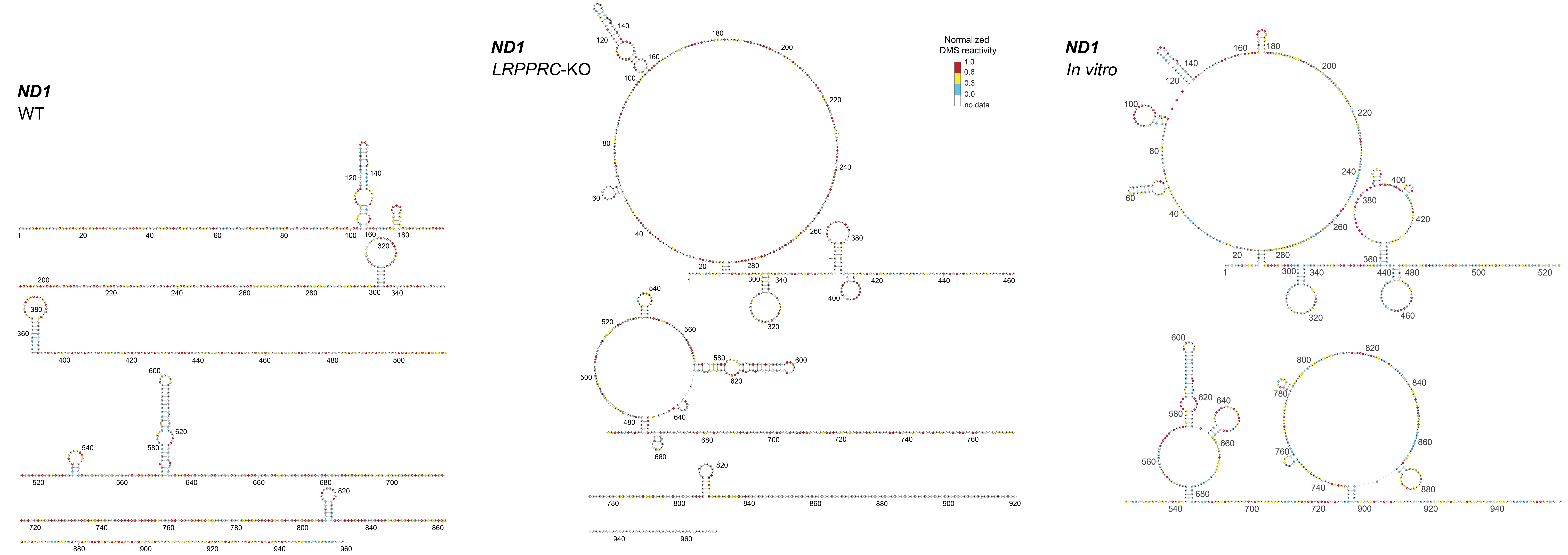

### Supplemental Datafile 5

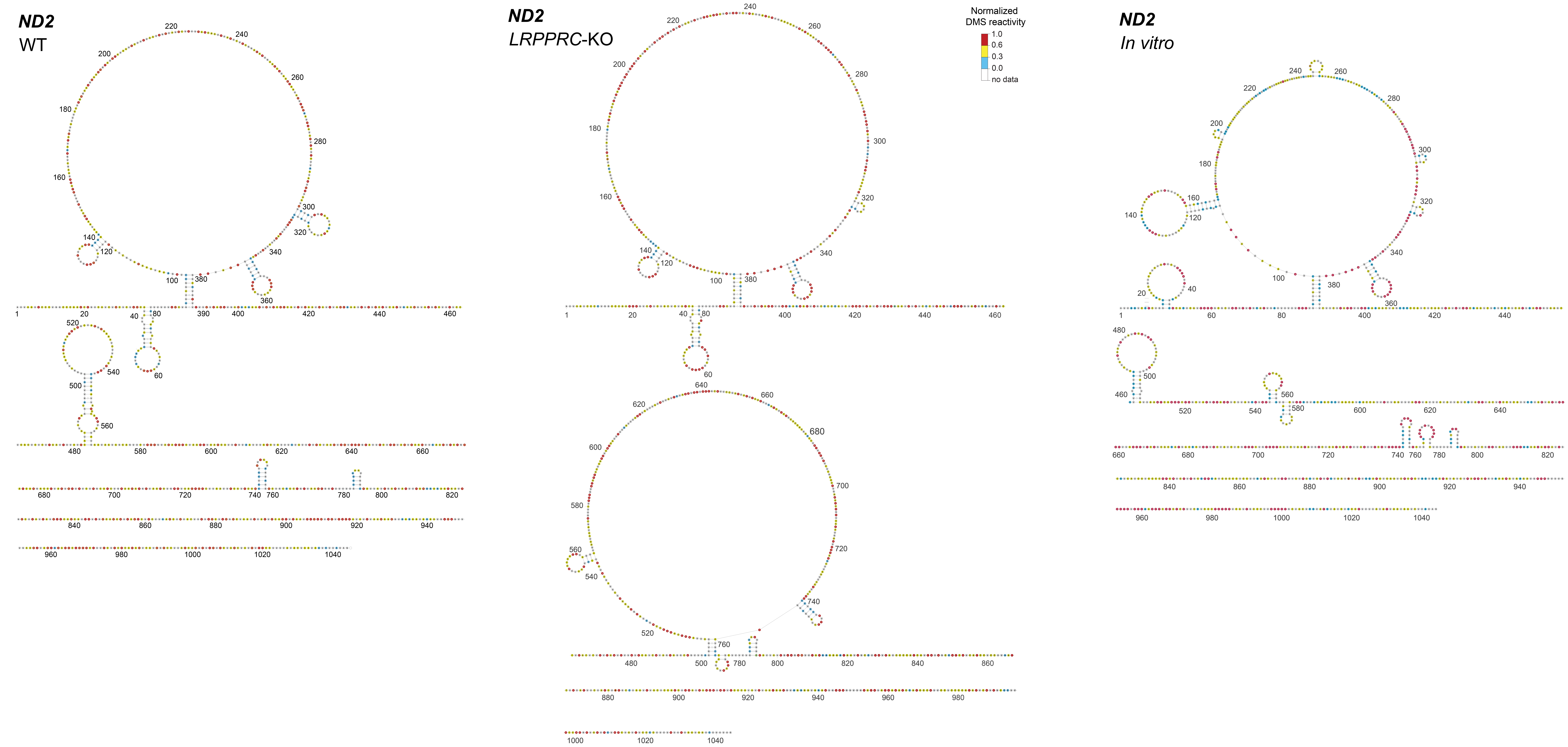

### Supplemental Datafile 6

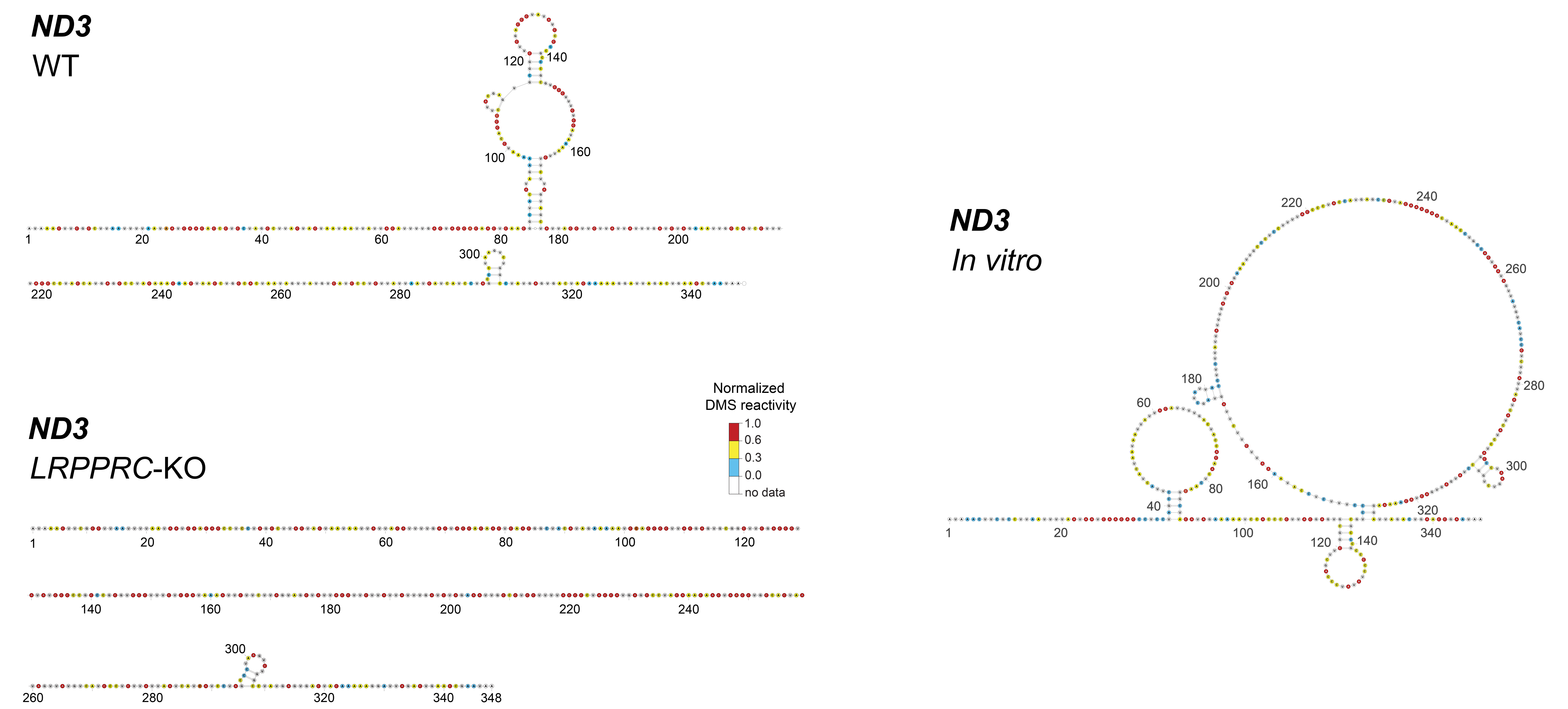

### Supplemental Datafile 7

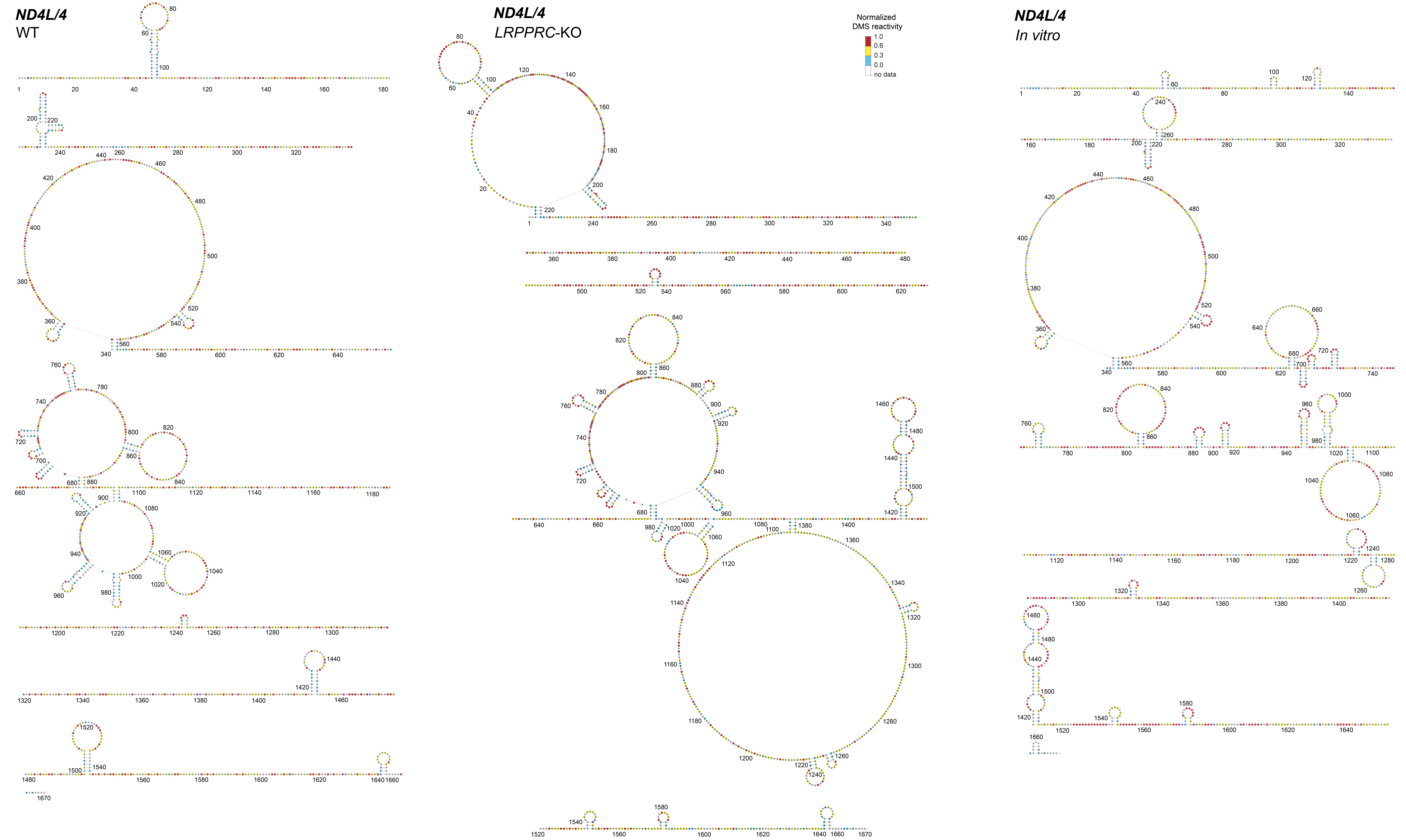

### Supplemental Datafile 8

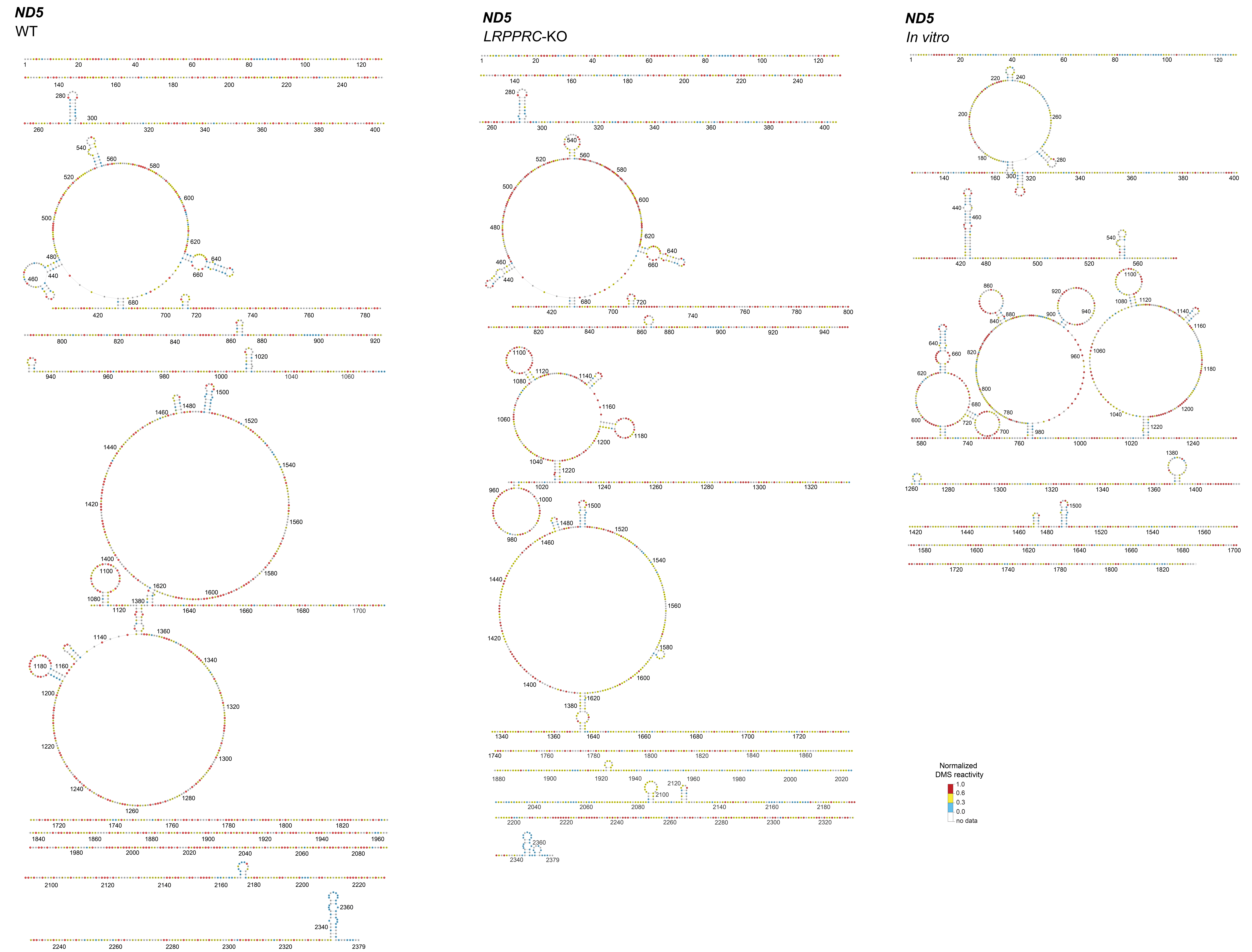

### Supplemental Datafile 9

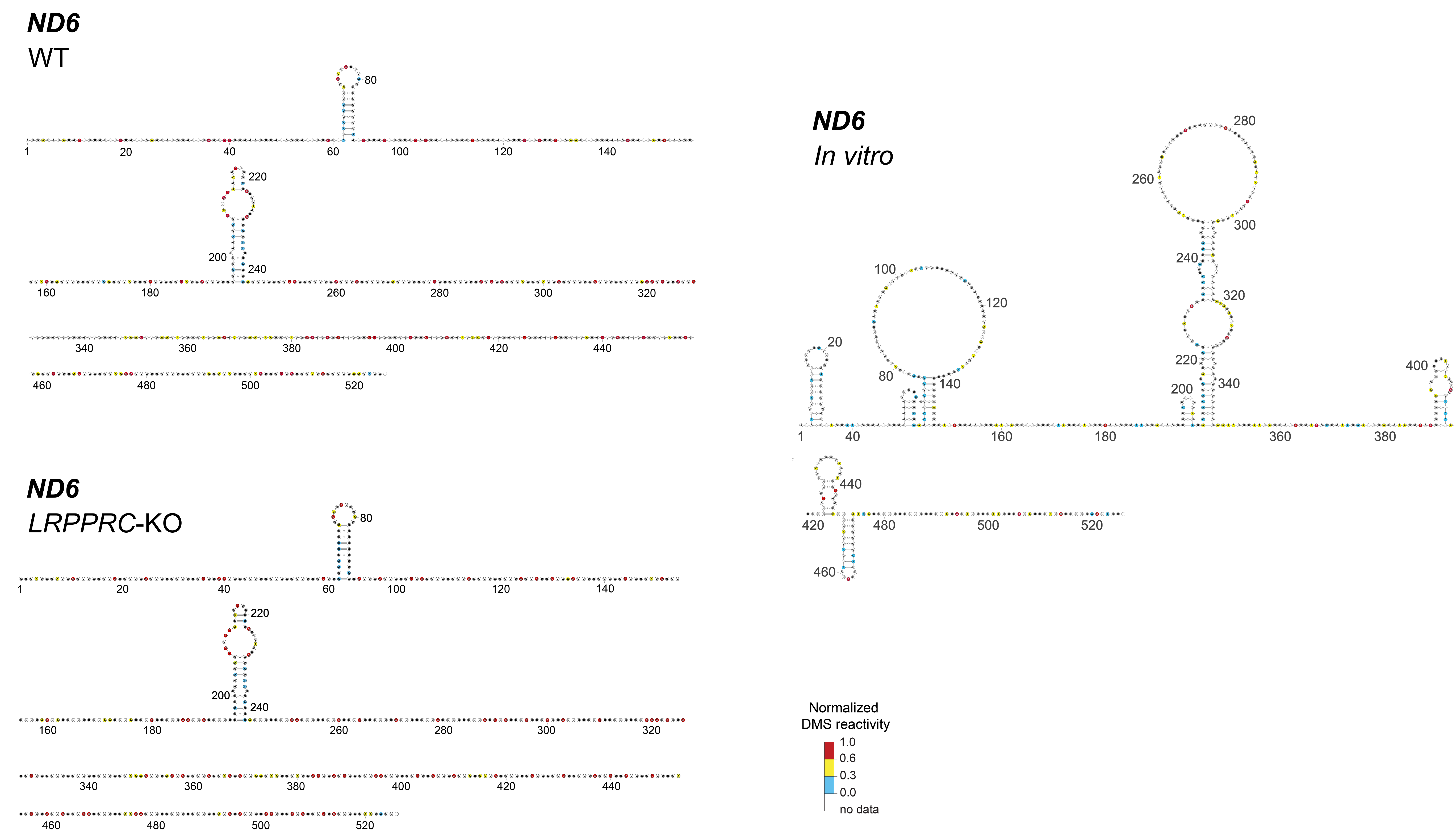

### Supplemental Datafile 10

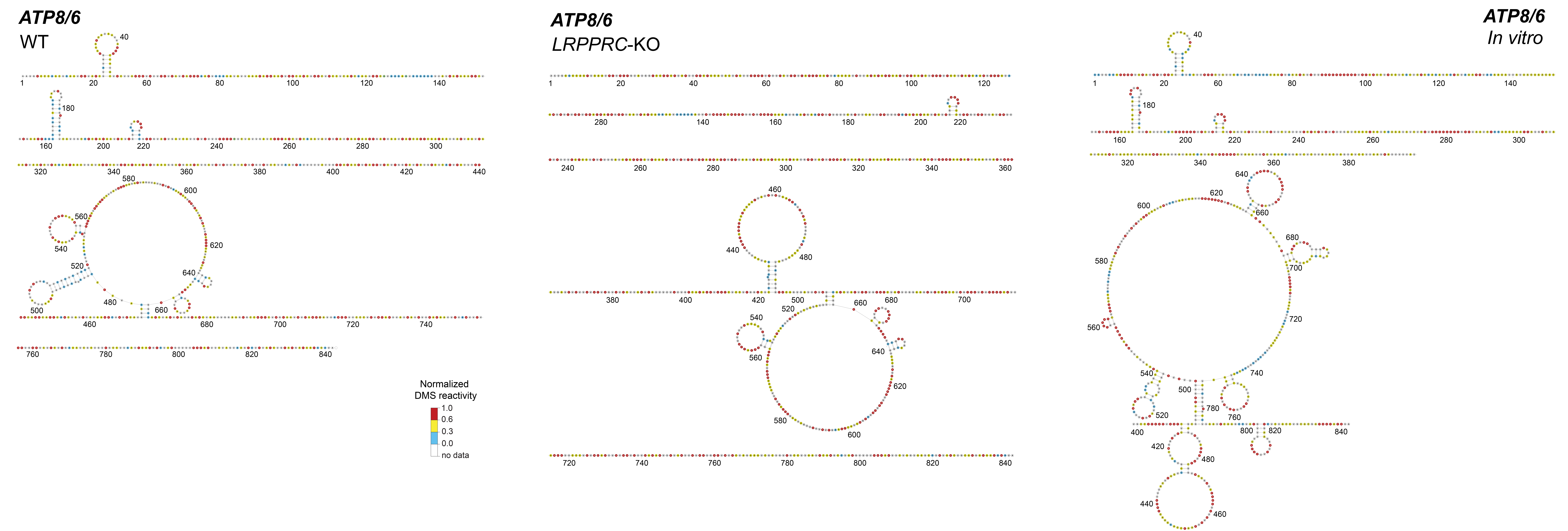

### Supplemental Datafile 11

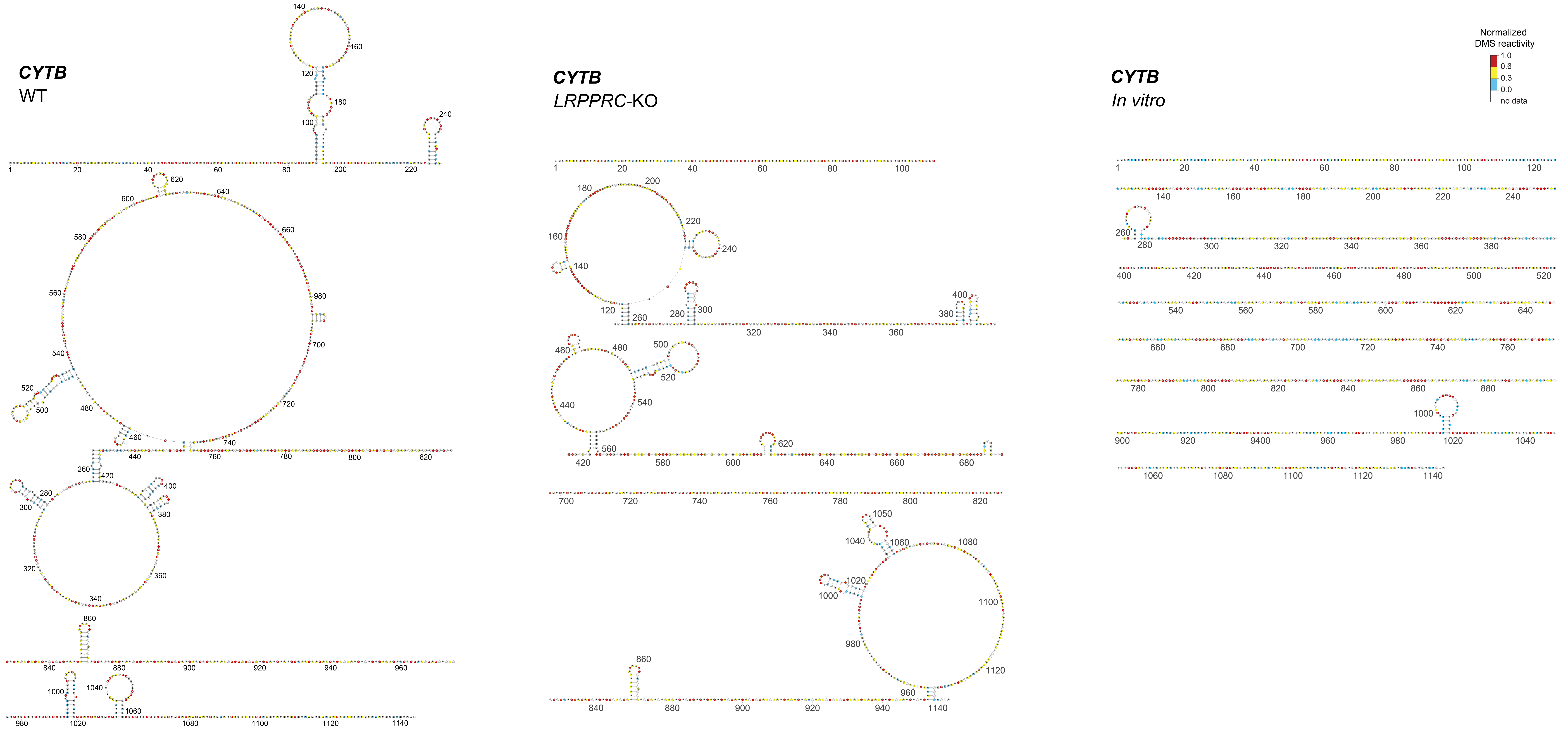
